# Differentiation reveals the plasticity of age-related change in murine muscle progenitors

**DOI:** 10.1101/2020.03.05.979112

**Authors:** Jacob C. Kimmel, David G. Hendrickson, David R. Kelley

## Abstract

Skeletal muscle experiences a decline in lean mass and regenerative potential with age, in part due to intrinsic changes in progenitor cells. However, it remains unclear if age-related changes in progenitors persist across a differentiation trajectory or if new age-related changes manifest in differentiated cells. To investigate this possibility, we performed single cell RNA-seq on muscle mononuclear cells from young and aged mice and profiled muscle stem cells (MuSCs) and fibro/adipose progenitors (FAPs) after differentiation. Differentiation increased the magnitude of age-related change in MuSCs and FAPs, but also masked a subset of age-related changes present in progenitors. Using a dynamical systems approach and RNA velocity, we found that aged MuSCs follow the same differentiation trajectory as young cells, but stall in differentiation near a commitment decision. Our results suggest that age-related changes are plastic across differentiation trajectories and that fate commitment decisions are delayed in aged myogenic cells.

## 1 Introduction

Aging entails a progressive functional decline over the lifespan of an organism, eventually resulting in death [40]. In mammalian muscle tissue, aging results in a decrease in muscle mass and regenerative potential. Loss of regenerative function has been attributed both to cell-intrinsic changes in muscle stem cells (MuSCs) and cell-extrinsic changes in the tissue environment [16, 12, 25]. Additional evidence suggests that age-related changes in fibro/adipose progenitors (FAPs), a mesenchymal progenitor population, may also play a role [12, 50]. Cell-intrinsic progenitor defects have been linked to reduced production of daughter cells and limited engraftment potential in aged MuSCs [17, 13, 28, 19]. However, it remains unknown if cell-intrinsic changes in aged progenitors are inherited by differentiated daughter cells, or if the differentiated progeny of these progenitors display unique age-related changes. It is also unclear if the trajectory, cell fates, and rate of differentiation are preserved with age in MuSCs and FAPs. Each of these possibilities suggests another route by which age-related changes in progenitors may manifest tissue-level defects.

During muscle regeneration, a subset of both MuSCs and FAPs shift cell identity and commit to cell fates by differentiation [14, 92]. Recent investigations of murine aging with single cell transcriptomics report that aging manifests uniquely across distinct cell identities [2, 44]. The relationship between aging and cellular differentiation however has been less explored. Cell identity changes during differentiation may alter the manifestation of aging, hiding differences apparent in the progenitors and uncovering new ones in the differentiated daughters.

Despite the marked functional defects of aged MuSCs during regeneration, transcriptional profiling studies have reported subtle age-related changes in MuSCs [51, 19, 77, 42]. Transcriptional age-related changes are similarly subtle in unchallenged cells from multiple tissues in recent single cell transcriptomics atlas studies [44, 83]. This disparity between functional defects and transcriptional changes suggests that some latent age-related changes may only manifest when cells are functionally challenged. In support of this perspective, a study of aged T cells indicates that immunological stimulation may elicit age-related changes that are otherwise latent [55].

Differentiation asks progenitor cells to undergo a challenging transformation and may also shed light on which differentially expressed genes and pathways are most relevant to aging phenotypes. Differentiation may also reveal age-related changes in more complex gene expression programs, such as changes in differentiation trajectories, differentiation fates, or the rate of fate commitment. Investigating gene expression changes revealed by differentiation may suggest genes that underlie functional decline in progenitors and provide a conceptual understanding of how fate commitment programs change with age.

It is difficult to address these gaps in knowledge with traditional cell biology techniques that can measure only a handful of molecular and behavioral markers in individual cells [16, 8, 19]. By contrast, recent developments in single cell genomics allow for the simultaneous profiling of the entire transcriptome across thousands of single cells [86]. Single cell genomics studies have already revealed novel aspects of aging in multiple tissues, including changes in the heterogeneity and distribution of cellular states and gene expression levels [46, 24, 2, 44, 83]. In the context of muscle aging, single cell genomics allows us to simultaneously identify the differentiation state of a progenitor cell and measure transcriptional features that may change with age.

To investigate these possibilities, we applied single cell RNA-seq (scRNA-seq) to profile muscle mononuclear cells from young and aged mice, both immediately following isolation and after a differentiation challenge to MuSCs and FAPs. We found latent age-related changes that are only revealed upon differentiation in both progenitor cell lineages. Differentiation also revealed that aged myogenic cells execute the differentiation program more slowly, stalling at a specific commitment decision, but preserve a youthful differentiation fate and trajectory. Our results indicate that the functional challenge of differentiation can reveal age-related changes at the level of individual genes and complex differentiation programs that would otherwise go unobserved.

## 2 Results

### 2.1 Single cell RNA-sequencing of muscle mononuclear cells identifies diverse cell types

We generated single cell RNA-seq profiles for muscle mononuclear cells from young (3 months old, *n* = 2) and aged (20 months old, *n* = 2) male C57Bl/6J mice. We immediately profiled a set of cells using the 10X Chromium microfluidics system (freshly-isolated timepoint). We also differentiated a set of young and aged MuSCs and FAPs from this isolation *in vitro* for 60 hours (differentiating timepoint) before transcriptional profiling (Fig 1A; Fig. S1; Methods).

**Figure 1:**
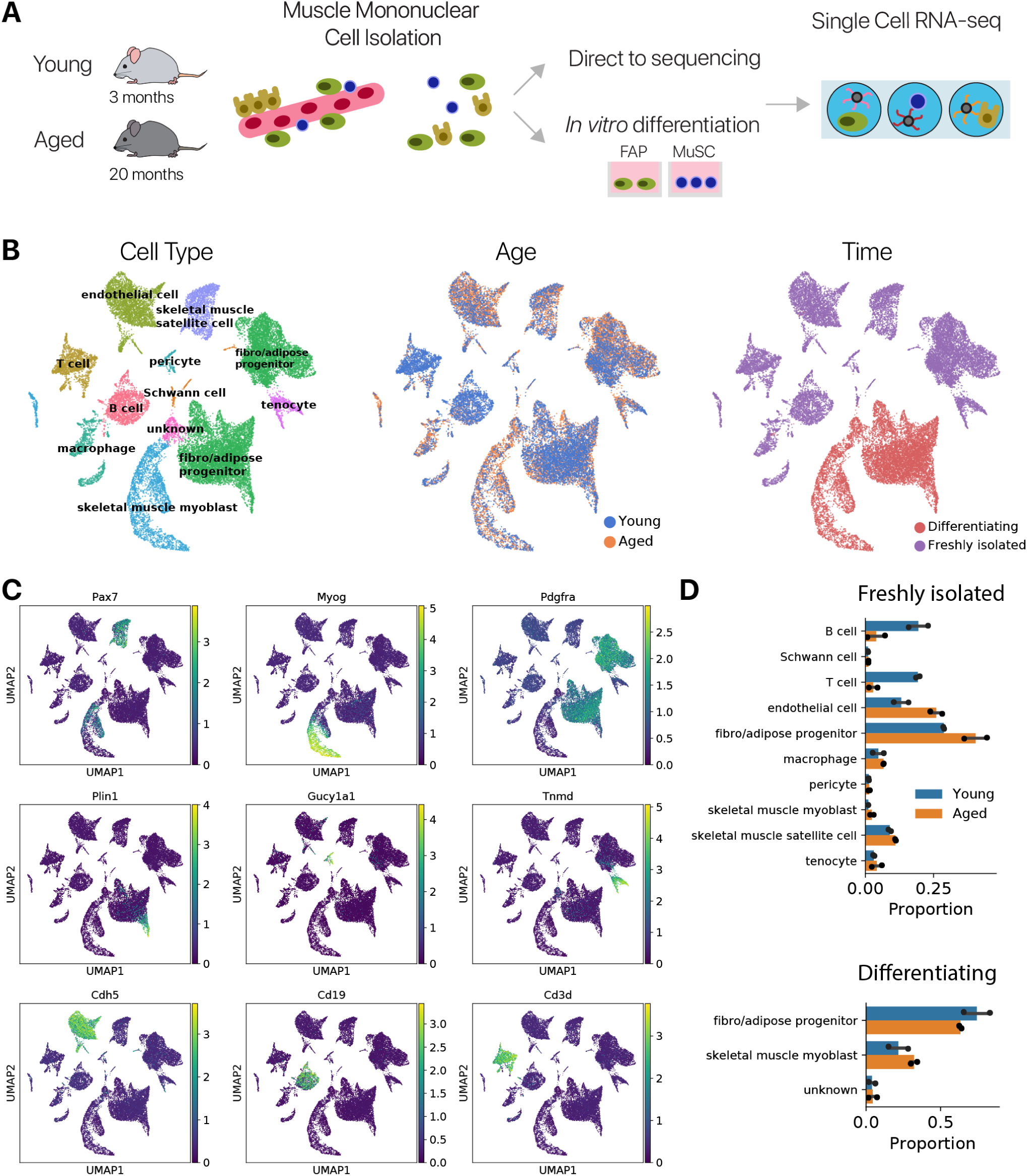
Single cell RNA-seq identifies diverse cell types in murine muscle mononuclear cells. **(A)** Experimental schematic. Muscle mononuclear cells were isolated from young and aged C57Bl/6 mice. Transcriptional profiles were collected immediately for a subset of cells. MuSCSs and FAPs were plated and allowed to differentiate for 60h before profiling. **(B)** UMAP projection of transcriptional space. Overlaid colors indicate identified cell types, the animal age, and the collection time point respectively. **(C)** Expression of cell type specific marker genes overlaid on UMAP projections. **(D)** Quantification of cell type proportions recovered from animals at each age and each timepoint. Freshly-isolated and Differentiating cells were profiled in different library preparations, such that relative proportions are not comparable between the timepoints.

After normalization and dimensionality reduction, we used a deep neural network classifier and manual validation to identify cell types from each relative mRNA abundance profile [44]. We identified skeletal muscle satellite cells (MuSCs), fibro/adipose progenitors (FAPs), endothelial cells, and multiple lymphoid cell types (Fig. 1B). Each of these cell identity annotations is supported by specific marker genes (Fig. 1C). We also identified pericytes marked by *Cspg4* and *Pdgfrb*, Schwann cells marked by *Plp1*, and tenocytes marked by *Tnmd* in our data. Although these cell types are not annotated in the *Tabula Muris* cell atlas, we found that these populations are present in the *Tabula Muris* data (Fig. S2). Pericytes are likewise unannotated in multiple single cell genomics studies of the limb muscle [21, 29, 20, 63]. We believe this is the first identification of this cell type in the muscle mononuclear population by single cell genomics, in agreement with histological literature noting its presence and importance [10, 11]. We found that *Gucy1a1* is a marker of this pericyte population (Fig. 1C). We recovered all cell types in all animals (Fig. S3).

Comparing across ages, we found that aged and young cells of each cell type cluster together and do not occupy distinct regions of a transcriptional latent space (Fig. 1B). This suggests that age-related changes are a smaller source of variation than cell type differences, as in previous single cell genomics studies of aging [24, 2, 44, 83]. We recovered comparatively fewer immune cells in aged animals at the freshly-isolated time point. However, we refrain from concluding this is biologically relevant because our FACS enrichment strategy may interact with the single cell dissociation efficiency with age (Fig. 1D, Methods).

Unique marker gene expression indicates that our differentiation treatment successfully induced a cell identity transition from muscle stem cells into differentiated myoblasts and myocytes [14]. We found that the late-stage myogenic regulatory transcription factor (MRF) *Myog* and sarcomere components *Myh1, Ttn*, and *Tnnc1* were uniquely expressed in cells from our differentiating time point (Fig. 1C, Fig. S4). FAPs have been reported to differentiate into adipocytes and myofibroblasts [39], with later studies reporting a myogenic *Peg3+* sub-population [58, 61]. Differentiating FAPs uniquely expressed adipocyte markers *Plin1* (Fig. 1C) and *Plin2*, and showed higher levels of myofibroblast marker *Acta2* (Fig. S4), confirming that our differentiation assay successfully induced FAPs to adopt multiple differentiated fates. To confirm that we recapitulate previously observed age-related changes, we examined the expression of aging marker genes in myogenic cells. Our data recapitulate some previously reported [51, 3, 42] age-related changes (Fig. S5), consistent with previous cross-study comparisons [42].

### 2.2 Differential expression identifies cell type specific and common age-related changes

Aging may manifest transcriptional differences within each cell type. To identify these age-related changes, we performed differential expression analysis between aged and young cells of each cell type at each timepoint (Methods). We first focused our analysis on cells that were profiled immediately following isolation as the best approximation of *in vivo* expression differences with age.

#### 2.2.1 Aged muscle stem cells lose quiescence and downregulate the unfolded protein response

In MuSCs, Gene Ontology enrichment analysis revealed that the unfolded protein response (UPR) was downregulated in aged cells (Fig. 2A, B). Previous studies have reported that the UPR is upregulated in quiescent MuSCs and helps maintain the quiescent stem cell state [97]. It has also been reported that aged MuSCs lose quiescence, suggesting the quiescence-associated UPR may be downregulated with age [16]. These previous observations suggest that the downregulation of the UPR with age that we observe here may be the result of aged MuSCs transitioning from a quiescent to a “primed” state [67, 68].

**Figure 2:**
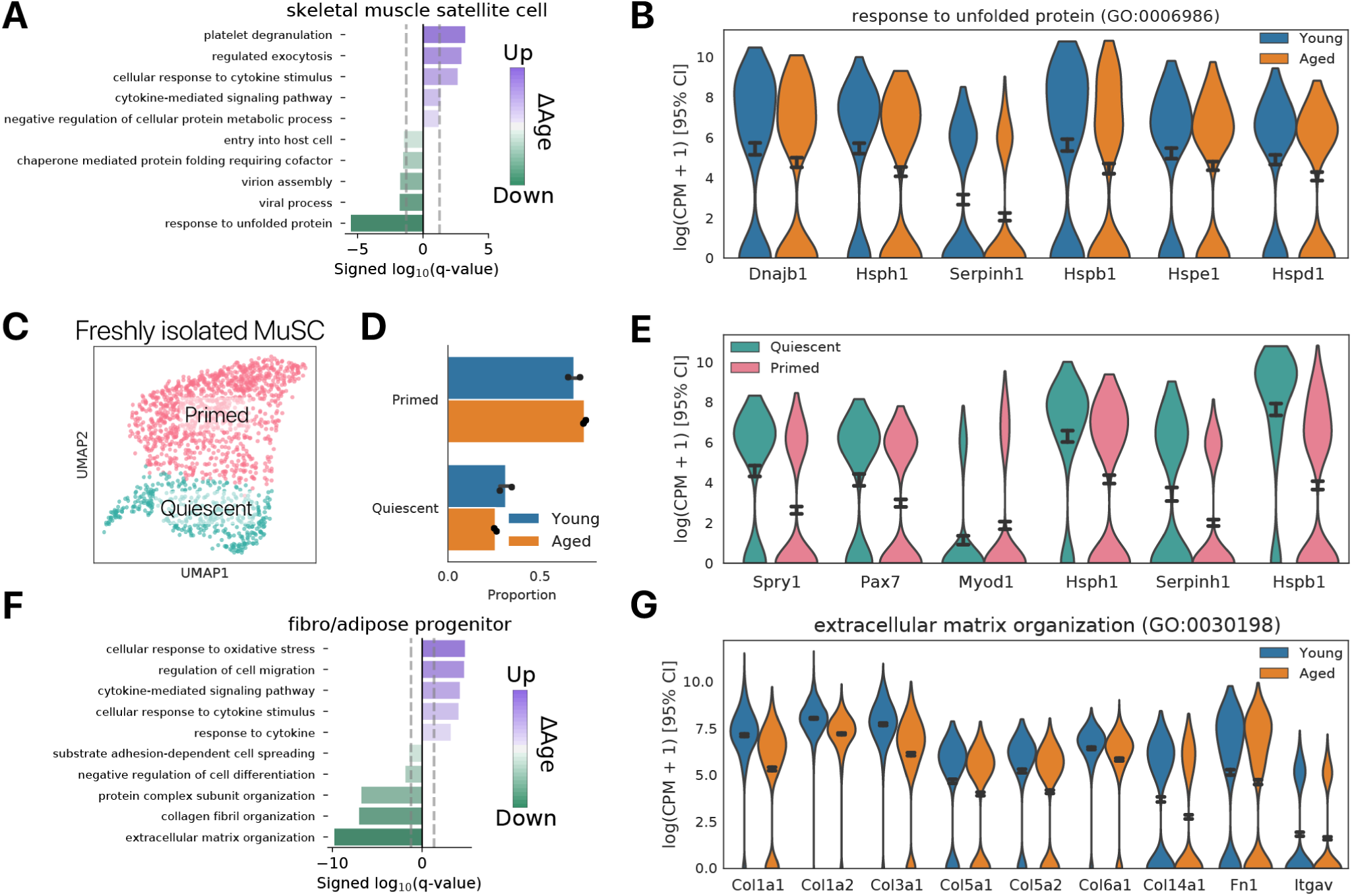
Differential expression reveals cell type specific and common aging phenotypes. **(A)** Gene Ontology enrichment analysis for differentially expressed genes between young and aged freshly-isolated muscle stem cells. **(B)** Violin plots showing the expression of a subset of genes driving enrichment of the “response to unfolded protein” GO term. **(C)** Leiden community detection identified two clusters of MuSCs in the freshly-isolated timepoint. One cluster represents “primed” MuSCs, enriched for *Myod1*, while the other captures quiescent MuSCs, enriched for *Spry1*. **(D)** Proportion of young and aged MuSCs in the primed and quiescent clusters. The primed cluster is enriched for aged MuSCs, while the quiescent cluster is enriched for young cells (*χ*^2^ test *p* < 0.01). **(E)** Expression of myogenic and UPR-associated genes across the quiescent and primed clusters. UPR genes are elevated in the quiescent cluster relative to the primed cluster. **(F)** Gene ontology enrichment analysis for differentially expressed genes between aged and young quiescent FAPs. **(H)** Violin plots showing the expression of a subset of genes driving enrichment of the “extracellular matrix organization” GO term.

To interrogate this hypothesis, we clustered the freshly-isolated MuSC population using Leiden community detection [85]. We found two populations corresponding to quiescent and primed MuSCs (Fig. 2C). The quiescent population was enriched for *Pax7* and for the quiescence-associated RTK inhibitor *Spry1* [75]. By contrast, the primed population was enriched for the canonical myogenic activation marker *Myod1* (Fig. 2E). Expression of UPR-associated genes was higher in the quiescent MuSC cluster, consistent with previous observations of high UPR activity in quiescent MuSCs (Fig. 2E). We also observed increased expression of cytokine response genes in aged MuSCs, suggesting immunological stress (Fig. S6B, C).

We found a modest enrichment of young cells in the quiescent MuSC cluster (*χ*^2^-test, *p* < 0.01), such that changes in the proportion of quiescent and primed cells may contribute to the age-related change in UPR activity (Fig. 2D). UPR-associated gene expression were also downregulated with age within each MuSC cluster, suggesting that differences in UPR activity were not purely the result of increases in the proportion of primed MuSCs with age (Fig. S6A). These results support prior reports of elevated UPR activity in quiescent MuSCs, and suggest that this UPR regulation may be disrupted during aging.

#### 2.2.2 Aged fibro/adipose progenitors and tenocytes downregulate extra-cellular matrix genes

We also performed Gene Ontology enrichment analysis on FAPs, revealing downregulation of multiple extra-cellular matrix (ECM) related gene sets (Fig. 2F). Multiple collagen genes showed decreased expression with age, indicating that collagen deposition from aged FAPs may be decreased (Fig. 2G). Collagen genes from multiple classes were downregulated (I: *Col1a1, Col1a2*, III: *Col3a1*, V: *Col5a1*), suggesting a broad decrease in collagen expression.

Gene Ontology analysis similarly revealed that tenocytes downregulated collagen genes with age (Fig. S8). Aged tenocytes showed decreased expression for multiple collagen types, again suggesting a general downregulation of collagen pathways rather than a shift between collagen types. We found that a subpopulation of cells expressing high levels of collagen was enriched for young cells (*χ*^2^-test, *p* < 0.01). Therefore, a shift in tenocyte cell states with age may contribute to changes in collagen expression (Fig. S8). Computing the change in collagen gene set activity with age across the nearest neighbor graph of observed cells further supports that FAPs and tenocytes are the dominant sources of change in ECM expression with age (Fig. S8H, Methods).

Age-related changes in ECM composition have previously been observed in the muscle [79, 27], with decreased collagen expression noted in bulk transcriptomics studies of aged rats [62]. However, the cellular origin of these changes remains unknown. These results suggest that collagen gene expression changes originate in FAPs and tenocytes, potentially leading to changes in ECM composition.

#### 2.2.3 Differential expression reveals common changes in protein localization and translation with age

In a previous study, we found that most age-related changes are cell type specific, with only a small set of common age-related changes [44]. Here, we found a small set of genes that are commonly (*k* >= 3 cell types) changed with age across cell types (Fig. S7A,B). Gene Ontology enrichment analysis revealed that SRP-dependent protein localization and co-translational protein targeting were commonly changed with age, as previously observed in the kidney, lung, and spleen (Fig. S7C). However, here we found that some components associated with these processes are upregulated while many more were down-regulated, suggesting that changes in these biological processes are not necessarily unidirectional among all players involved.

### 2.3 Differentiation exacerbates the magnitude of age-related change in MuSCs and FAPs

Differentiation may influence the magnitude of age-related change, increasing or decreasing the distinctiveness of young and aged cell populations. To measure the magnitude of aging in a rigorous manner, we computed maximum mean discrepancy (MMD) distances. The MMD distance provides an estimate of the difference between two samples of cells, capturing differences in means, covariance, and modality [30]. We computed MMD distances between young and aged cells of each type using log-normalized counts as input features to estimate the magnitude of aging (Methods).

We found that all cell types at both timepoints exhibited a significant magnitude of aging (Kernel Two-Sample test, *p* < 0.001), while isochronic negative control comparisons showed non-significant MMD values near zero as expected (Fig. S9A, B; mean *p* > 0.99). Myogenic cells and FAPs displayed a higher magnitude of aging after differentiation, consistent with higher classification accuracy after differentiation (Wilcoxon rank sums, *p* < 0.001, Fig. S9A). Macrophages and FAPs show the highest MMD in the freshly-isolated time point, suggesting these cell types exhibit the largest differences with age in the unchallenged, *in vivo* context. This result further suggests that the cell identity changes induced by differentiation can reveal age-related change unseen in the unchallenged context.

Significant MMDs for each cell type confirm that aging is a substantial perturbation to each cell population, even if it is not the largest source of variation in our data. We also trained age classification models to estimate the distinctiveness of cell populations, and these models produced similar results (Fig. S10).

### 2.4 Trajectories and cell fates of myogenic differentiation are preserved with age

Aging may alter cellular differentiation trajectories and fates, as observed in hematopoiesis [94] and suggested for muscle [13, 70]. We employed pseudotime analysis to infer a trajectory of myogenic differentiation in both young and aged cells to investigate this possibility. Prior to identifying a pseudotime trajectory, we removed cycling and apoptotic myogenic cells *in silico* (Fig. S11, Methods). We found a smooth trajectory of cells with markers of myogenic progenitors (*Pax7*) at one end and commitment markers at the other (*Myog, Myh1*; Fig. 3A, S12A).

**Figure 3:**
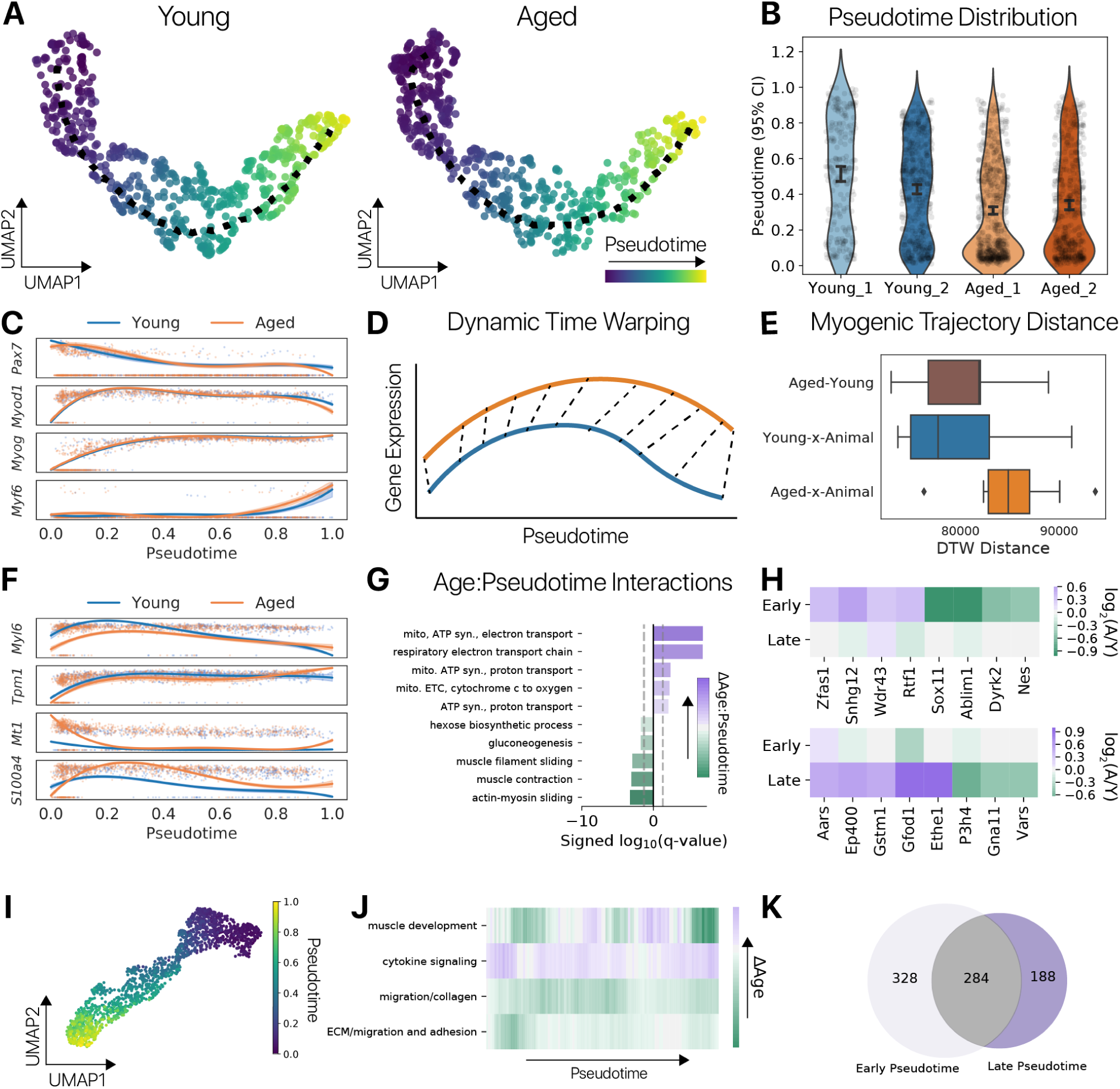
Myogenic differentiation masks and reveals age-related changes. **(A)** Pseudotime trajectories inferred for young and aged cells independently. The black line represents the mean coordinate in transcriptional space across the trajectory. **(B)** Distribution of myogenic cells along a common pseudotime coordinate, displayed for each animal. Young cells are significantly enriched at the end of the pseudotime axis (ANOVA, *p* < 0.001). **(C)** Gene expression over pseudotime for *Pax7* and myogenic regulatory factors. Curves are spline fits and points are expression in individual cells. Values are scaled expression ([0, 1]) and shading represents ± SD. Kinetics of myogenic gene expression are qualitatively preserved across ages. **(D)** Cartoon illustration of dynamic time warping (DTW) across pseudotime axes. DTW distance is computed between young and aged spline fits for each gene. **(E)** DTW distances for heterochronic (Aged-Young) pseudotime trajectories compared to distances computed between animals of the same age (Aged-x-Animal, Young-x-Animal). **(F)** Gene expression over pseudotime for genes with a significant Age:Pseudotime interaction (*q* < 0.05, likelihood-ratio test). Some genes show amelioration of age-related differences over pseudotime, while others show increases in age-related change. **(G)** Gene Ontology analysis of genes with significant Age:Pseudotime interactions revealed that mitochondrial respiration genes have a positive interaction across differentiation, while muscle contraction genes have a negative interaction. **(H)** Differential expression (log_2_Aged/Young) of genes that are “masked” (upper) and “revealed” (lower) by differentiation. All genes shown are significantly differentially expressed at one end of the pseudotime trajectory (Early, Late), but not the other. All genes are expressed at both ends of the trajectory. **(I)** NMF embedding of differentiating myogenic cells projected with UMAP. **(J)** Age-related changes in selected NMF-derived gene expression programs over pseudotime. Some expression programs show larger differences with differentiation (muscle development), while others expression programs are normalized (ECM/migration). **(K)** Venn diagram showing the overlap of differentially expressed gene sets in early and late stages of the pseudotime trajectory.

We used diffusion pseudotime analysis [33] to assign a pseudotime coordinate to each cell from young and old animals, given a root cell chosen from *Pax7+* progenitors (Fig. 3A, Methods). We found that young cells were significantly advanced in pseudotime relative to aged cells (ANOVA *p* < 0.001, Fig. 3B). The difference was insensitive to the selection of root cell (Fig. S12B). This result suggests that young cells progress more rapidly through differentiation than aged cells. This observation stands in contrast with some reports that have suggested that aged myogenic cells may experience “precocious” commitment [8, 16]. As our results are consistent with another recent transcriptomic analysis showing aged MuSCs express higher levels of quiescence marker *Pax7* [63], the contrast with previous reports may be explained by the reliance of previous studies on marker gene analysis within subsets of the myogenic cell population.

To determine if aging changes the dynamics of gene expression during differentiation, we fit quartic splines to the expression of each gene over pseudotime for young and aged cells. We performed differential expression analysis using nested hurdle models (Methods) to identify genes with significant Age:Pseudotime interactions and detected only 91 genes with dynamics altered by age. We investigate these genes in the next section. The stem cell marker *Pax7* and the myogenic regulatory factors displayed similar behavior across pseudotime in young and aged cells (Fig. 3C, *q* > 0.05 log-likelihood ratio test, Age:Pseudotime interaction). These results suggest that the path of differentiation in gene expression space is preserved in aged myogenic cells.

To determine if aging alters the shape of the commitment trajectory when considering all genes, we inferred separate pseudotime trajectories for young and aged cells. We compared the difference between young and aged trajectories using a dynamic time warping (DTW) distance between splines fit to the expression of each gene across pseudotime, as previously described [15] (Methods). This process is illustrated for a single gene in (Fig. 3D). To account for variation introduced by the selection of a root cell and random sampling of cells during scRNA-seq, we randomly sampled subsets of cells from each age, chose a random root cell from the *Pax7+* population, and computed DTW distances across these bootstrap samples (Methods).

We measured distances between young and aged trajectories in a heterochronic comparison, and also performed isochronic young to young and aged to aged comparisons across animals to estimate a negative control distribution. We found that the trajectory distances between isochronic cells from different animals are comparable to our heterochronic comparison (Fig. 3F). Therefore, the heterochronic trajectory distance we observed is on par with the expected distance between trajectories observed from different animals of the same age. This result suggests that the trajectory of myogenic differentiation is not significantly different between young and aged mice. Rather, the majority of gene expression changes that occur with fate commitment are preserved with age.

Consistent with the preservation of the myogenic differentiation trajectory, we found no evidence for aberrant, fibrogenic cell fates [13, 70]. Both young and aged MuSCs exhibit a common myocyte fate, marked by expression of *Myog* and *Myh1* (Fig. S12A) [14]. We found that no collagen genes were differentially expressed with age, and the activity of the Gene Ontology collagen fibril organization gene set was marginally higher in young MuSCs (Fig. S12C). Both of these results are inconsistent with an alternative, fibroblast fate. These data therefore suggest that both the trajectory and fate of myogenic differentiation are preserved in aged MuSCs.

### 2.5 Myogenic differentiation reveals age-related gene expression changes

We identified 91 genes with significant Age:Pseudotime interactions, suggesting that differentiation influences age-related change in myogenic cells. Within the significant Age:Pseudotime interaction genes, we found examples of genes whose magnitude of age-related change increased with differentiation (*Ttn*), decreased with differentiation (*Mt1*), or displayed non-monotonic behavior (*S100a4*, Fig. 3F). Multiple genes with Age:Pseudotime interactions were expressed across the entire trajectory, indicating the interaction was not a simple on/off switch with differentiation (e.g. *Myl6, S100a4*).

Performing Gene Ontology analysis, we found that genes with positive Age:Pseudotime interactions were enriched for mitochondrial gene sets, while negative interactions were enriched for myogenic differentiation gene sets (Fig. 3G). Association of age with higher mitochondrial oxidative phosphorylation is consistent with aged myogenic cells failing to adopt differentiated myogenic states [69]. These results suggest that differentiation both masks and reveals age-related changes.

To further explore the influence of differentiation on age-related change, we binned myogenic cells into three groups based on their pseudotime coordinates and computed differential expression across ages within each bin. We considered the first of these bins to be primitive cells, “early” in pseudotime, and the last bin to be more committed cells, “late” in pseudotime. We found more than 300 genes that were changed with age in primitive (early) cells, but not in committed (late) cells, suggesting these age-related changes are masked by differentiation (Fig. 3; Fig. 3H, upper). The majority of these genes (303) are expressed in more than 100 cells in the both early and late populations, indicating that the masking of age-related change with differentiation is not a simple “on to off” switch. Conversely, we found 188 genes that were changed with age in committed (late) cells, but not in primitive (early) cells (Fig. 3H, lower). The majority of these genes (176) were expressed in > 100 early, primitive cells, suggesting that these age-related changes revealed by differentiation are not due to simple “off to on” switches.

We found similar results by comparing age-related changes in primitive and committed cell populations isolated based on marker genes *in silico* (Fig. S12A, B, C, D) and when comparing across timepoints in our experiment (Fig. S14E, F). Together, these analyses suggest that differentiation masks some age-related changes present in progenitors, while revealing age-related changes unseen before differentiation. These results cannot be explained by simple on/off switching of cell identity genes during differentiation, as many of the masked and revealed age-related changes occur in genes that expressed in both progenitors and differentiated cells.

To investigate changes in the activity of gene expression programs, we embedded differentiating cells with non-negative matrix factorization (NMF) to capture coordinated expression changes that involve many genes [80, 81, 45](Fig. 3I, Fig. S15, Methods). We derived semantic labels for these programs using gene set enrichments (Fig. S16, Methods) In the NMF embedding, we compute an “aging trajectory” as the distance between the centroid of young and aged cells, capturing differences in gene expression program activity with age [44](Methods). Here, we used a sliding window approach to compute an aging trajectory across pseudotime (Fig. 3J, Methods).

Similar to results at the gene level, we found that the aging trajectory shifts across the pseudotime trajectory. The degree of age-related change increased with differentiation in some gene programs (muscle development) and decreased in other programs (ECM/migration and adhesion). Other programs exhibited non-monotonic behavior across pseudotime, with the maximum age-related change observed at some point mid-way through differentiation (migration/collagen). These results are consistent with our gene level analysis and further suggest that differentiation masks and reveals age-related change.

#### 2.5.1 Fate-specific age-related changes are revealed by fibro/adipose progenitor differentiation

We similarly explored whether FAP differentiation masks and reveals age-related gene expression changes. Examining marker gene expression in differentiating FAPs, we found subpopulations marked by adipocyte markers (*Plin1, -2, -4*), myofibroblast markers (*Acta2*/*α*SMA, *Vim*), and a population of pre-adipocytes enriched for progenitor markers (*Cd34, Ly6a*) (Fig. S17F). We decomposed the differentiating FAPs into these distinct differentiation fates using Leiden clustering (Fig. 4A). We fit a pseudotime trajectory to the myofibroblast and adipocyte fates with diffusion pseudotime, using PAGA [90] to define unique paths for each fate (Methods, Fig. 4B). These pseudotime trajectories agreed with the directionality of RNA velocity estimates [47](Fig. S18). In contrast to myogenic cells, we observed no significant difference in the distribution of young and aged cells along these trajectories (Fig. S17H, I).

**Figure 4:**
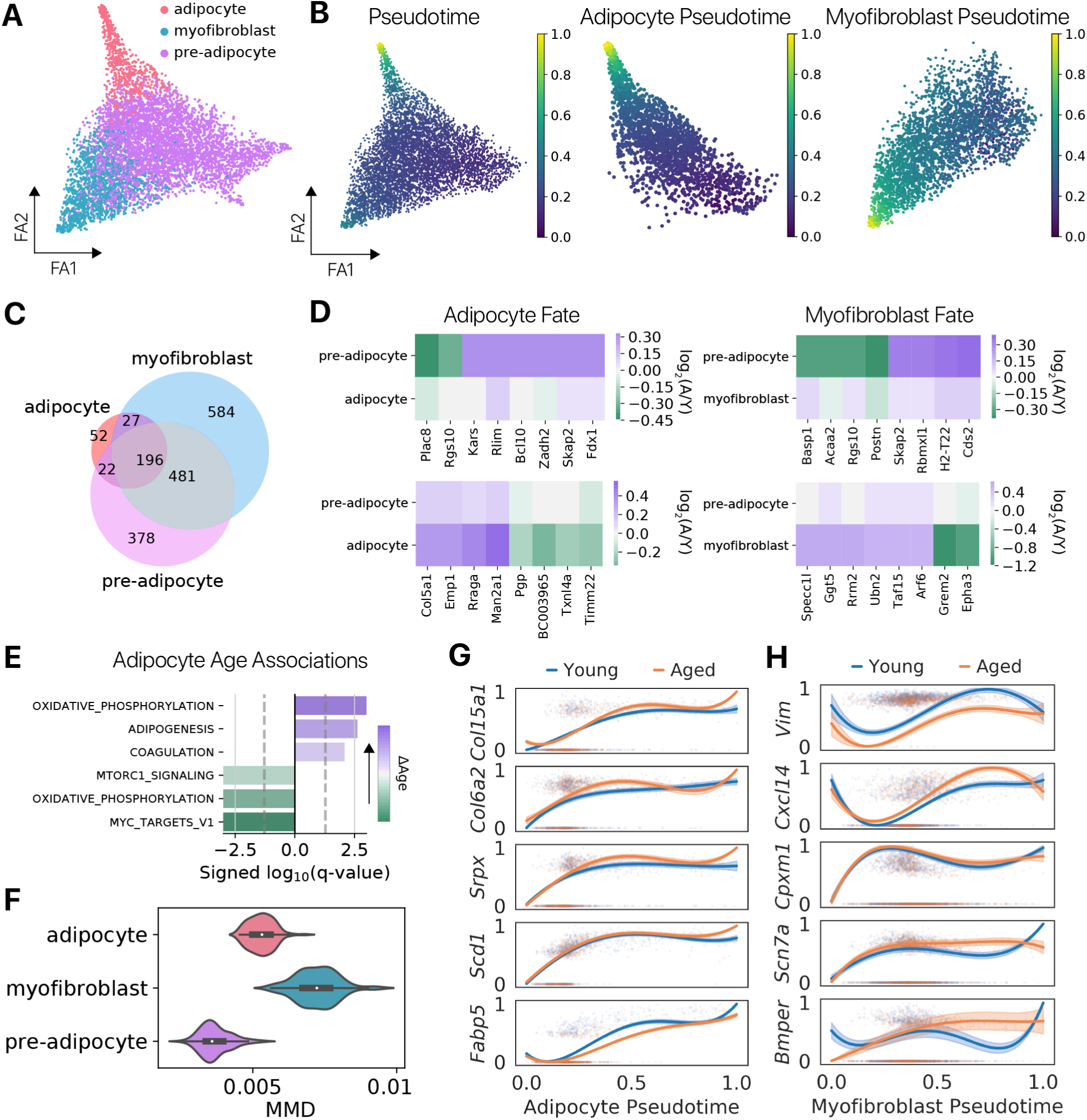
Age-related changes depend on FAP cell fate selection. **(A)** Force-directed graph layout (ForceAtlas2, FA2) for fibro/adipose progenitors (FAPs), colored by cell state annotations. **(B)** Diffusion pseudotime computed for the whole FAP population and pseudotime coordinates normalized across the adipocyte or myofibroblast differentiation path displayed on the graph layout. **(C)** Each FAP differentiation fate displays unique differentially expressed genes, with only a small set of genes shared between fates. **(D)** Differential expression (log_2_Aged/Young) of genes that are “masked” (upper) and “revealed” (lower) by differentiation in either the adipocyte or myofibroblast fate. All genes shown are expressed in both the progenitor and differentiated cell populations. **(E)** Genes upregulated with age in the adipocyte trajectory (*q* < 0.05, likelihood-ratio) are enriched for “adipogenesis” and “oxidative phosphorylation” gene sets (Hallmark, MSigDB, Fisher’s exact *q* < 0.05). **(F)** Maximum mean discrepancy distances computed between random samples of young and aged cells within each differentiation state. Both adipocytes and myofibroblats exhibit a larger magnitude of aging than pre-adipocyte progenitors. **(G)** Expression of genes with significant Age:Pseudotime interactions across adipocyte differentiation and **(H)** myofibroblast differentiation. Curves are splines, points are individual cells, shading represents ±SD. Age-related changes increase across pseudotime for some genes, and decrease with pseudotime for others.

We compared age-related transcriptional changes between the distinct differentiation fates (adipocytes, myofibroblasts, pre-adipocytes). We found that each fate exhibited unique age-related changes, with only a minority of differentially expressed genes shared between any two fates (Fig. 4C). A set of age-related changes in pre-adipocyte progenitor cells were masked after cells adopted an adipocyte or myofibroblast fate (Fig. 4D, upper). Conversely, there were age-related changes in adipocytes and myofibroblasts that were not observed in pre-adipocyte progenitors (Fig. 4D, lower). We similarly found age-related changes that were masked and revealed by differentiation when comparing across timepoints (Fig. S17A, B).

The majority of genes whose age-related change was masked or revealed by fate commitment were expression in both pre-adipocytes and the relevant cell fate, suggesting that masked and revealed changes are not the result of “on/off” switches in identity program genes (> 89% of genes are expressed in > 100 cells in each condition in each contrast). The set of genes that is masked and revealed by differentiation in this manner is unique for each fate. These results suggest that age-related change may be masked and revealed by cell differentiation in a fate-specific manner.

As in the myogenic population, we computed differential expression using nested hurdle models to identify significant Age:Pseudotime interactions for both the adipocyte and myofibroblast pseudotime trajectory (Methods). These analyses revealed 30 significant Age:Adipocyte Pseudotime interactions and six significant Age:Myofibroblast Pseudotime interactions (*q* < 0.05, log-likelihood ratio test). We also computed differentially expressed genes with age for all cells in the adipocyte and myofibroblast trajectories, using the pseudotime trajectory as a covariate. Genes upregulated with age in adipocytes are enriched for “adipogenesis” genes (Fig. 4E), while genes upregulated with age in myofibroblasts are enriched for “TGF*β* signaling” genes (Hallmark gene sets, MSigDB, *q* < 0.05, Fig. S17G). TGF*β* signaling is a canonical driver of fibroblast to myofibroblast conversion [88]. These results suggest that fate-specific age-related changes manifest during FAP differentiation.

Investigating individual Age:Adipocyte Pseudotime interactions, we found age-related changes increased with differentiation (*Col15a1, Scd1*) as well age-related changes that were largest midway through differentiation (*Fabp5*, Fig. 4G). We similarly found multiple relationships between age-related change and Myofibroblast Pseudotime (Fig. 4H). These results complement our coarse-grained analysis and provide evidence that age-related changes may be masked and revealed by differentiation.

To determine if FAP differentiation influences the magnitude of aging, we computed MMD distances between young and aged cells within each FAP differentiation state. Both adipocytes and myofibroblasts exhibit larger magnitudes of aging than pre-adipocyte progenitors, suggesting that differentiation reveals age-related change not evident in the progenitor population (Fig. 4F). Combined with our results in myogenic commitment, these results suggest that cell identity changes during cell differentiation can mask and reveal age-related change. Observation of this phenomena in two distinct cell lineages suggests it is not specific to a particular differentiation program. Rather, the masking and revelation of age-related change may be a general feature of cell identity changes that occur during differentiation.

### 2.6 RNA velocity reveals a energy barrier in myogenic differentiation

Our observation that young myogenic cells are enriched in the late portions of the pseudotime trajectory (Fig. 3C) suggests that young and aged cells may progress through myogenic differentiation at different rates. To investigate this hypothesis, we employed RNA velocity analysis to infer rates of transcriptional change [47]. RNA velocity uses the number of reads mapping to intronic regions of a transcript – indicating an unspliced, nascent transcriptional product – to estimate a rate of change for each gene. Recent work has reported that these inferences reflect experimentally measured RNA turnover rates [66]. A cell’s velocity profile across all genes reflects the estimated change in its transcriptional state. We embedded velocity profiles and display RNA velocity estimates as vectors on the PCA projection (Fig. 5A).

**Figure 5:**
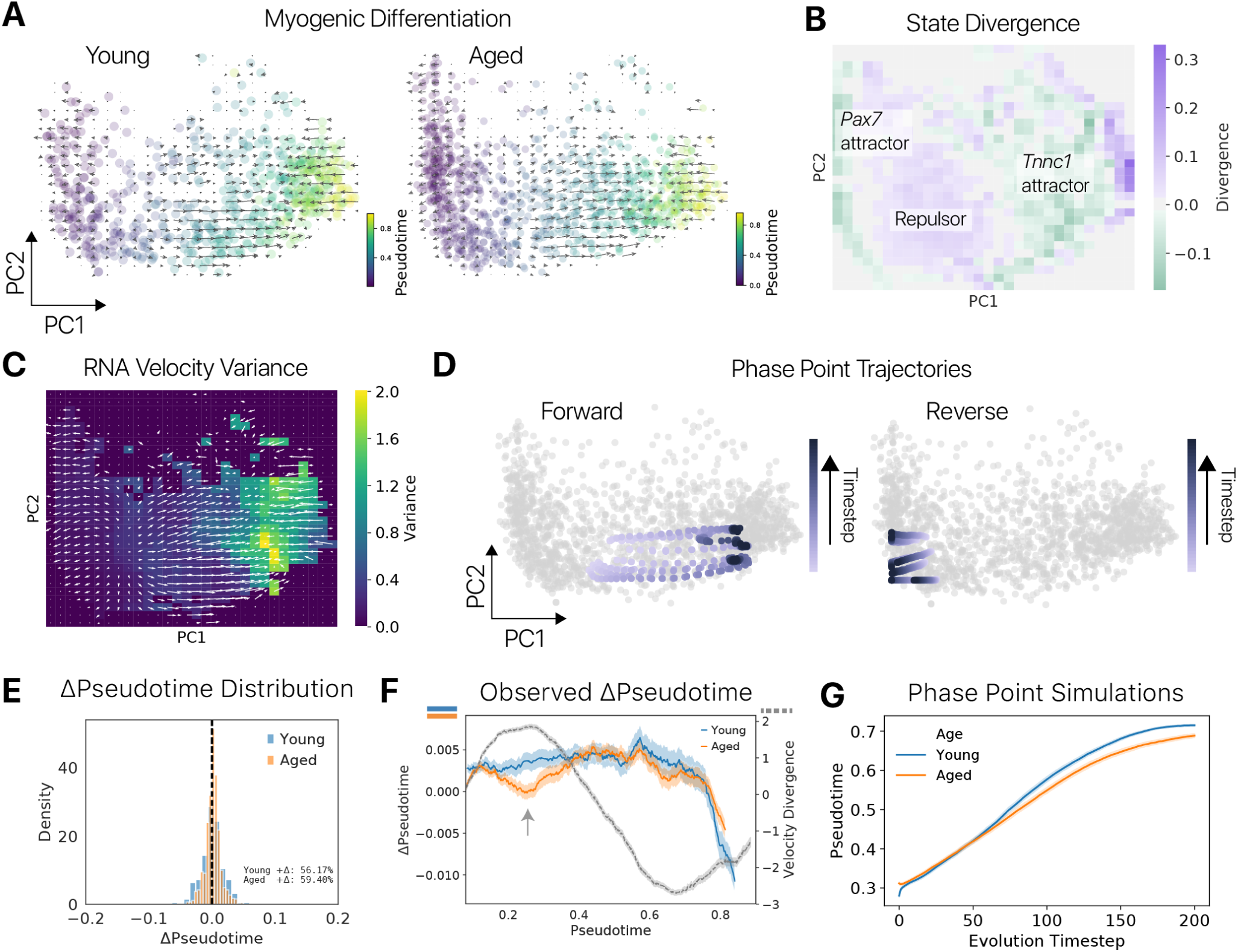
RNA velocity inference reveals an energy barrier between self-renewing and committed myogenic fates. **(A)** RNA velocity vectors atop separate PCA embeddings of young and aged cells. Velocity is qualitatively similar between young and aged cells. **(B)** State divergence, the relative “flow” of cells from a given region of state space, computed from RNA velocity vectors in a coarse-grained PCA embedding. We found two metastable attractor states (negative divergence, green) corresponding to “reserve cells” and differentiated cells. These attractors are separated by an unstable repulsor (positive divergence, purple) state that may represent a decision point for lineage commitment. **(C)** Variance of the RNA velocity vectors across the PCA embedding. Variance is computed on a grid of points using a Gaussian kernel to weight the contribution of nearby vectors. **(D)** Example trajectories from phase point simulations initiated on either side of the repulsor state shown atop the PCA embedding. Points initiated after the repulsor progress through the trajectory toward the *Tnnc1* committed state, while points initiated prior to the repulsor move toward the *Pax7* reserve cell state. **(E)** Normalized distribution of pseudotime changes predicted using RNA velocity for young and aged cells. Young cells display a broader distribution, with more probability mass on positive ΔPseudotime values. **(F)** Change in pseudotime predicted for cells across the pseudotime trajectory (rolling mean ± SEM). Aged cells show a lower proportion of cells moving forwards in pseudotime near the location of the repulsor state (pseudotime ≈ 0.25, OLS log-likelihood ratio test, *p* < 0.001). This region corresponds to the repulsor based on high velocity divergence (right axis, grey line). **(G)** Young phase points advance more rapidly than aged phase points when simulations are initiated after the repulsor state.

Comparing RNA velocity estimates in the embedding space between young and aged cells, we found few qualitative differences (Fig. 5A). We observed a bifurcation in the direction of velocity vectors in the early portion of the trajectory, roughly corresponding to the boundary between *Pax7* and *Myog* expression (Fig. S19F). This bifurcation may represent a decision point during the process of fate commitment.

To investigate this possibility, we computed the divergence of the RNA velocity field. The divergence reflects the net flow of cells to a particular region of state space and is frequently used to characterize vector fields in physical systems [82]. Negative divergence indicates a state is metastable, with more cells entering than exiting, while positive values indicate the opposite. We binned the space spanned by the first two dimensions of the PCA embedding and computed a weighted mean velocity for each bin in this discretized grid. From this discretized velocity, we computed a discretized divergence and performed bootstrapping to assess the significance of these divergence values (Methods).

We found that the bifurcation point in the velocity field represents an unstable repulsor state, with stable attractors on either side (Fig. 5B). There were minimal qualitative differences in the divergence of young and aged velocity fields (Fig. S21). In both cases, we found that the first attractor (earliest in pseudotime) is co-localized with expression of *Pax7*. Cells moving “backward” in pseudotime toward this *Pax7* high state may represent reserve cells that fail to differentiate or become more primitive in response to differentiation cues [93].

The second attractor is co-localized with expression of *Tnnc1*, a troponin component and marker of late-stage myogenic differentiation. Cells moving “forward” in pseudotime toward this attractor represent those that are committed and progressing through differentiation. Investigating the contributions of individual genes to RNA velocity vectors, we found induction of myogenic genes to be the primary source of forward velocity, while repression of myogenic genes and induction of non-myogenic genes was the primary source of backward velocity (Fig. S20, Methods).

We hypothesized that cells near the repulsor would show greater empirical variance in RNA velocity across cells, as different subsets move forward or backward. Instead, we found greater variance in the *Tnnc1* attractor, suggesting that cells in this basin move with random directionality in state space, rather than becoming stationary (Fig. 5C, Methods). Increased variance in the *Tnnc1* attractor accounts for the difference in ΔPseudotime variance between young and aged cells (Fig. S19B, E). This result is consistent with a recent correlation based metric of velocity uncertainty (Fig. S19F, Methods)[7].

To quantitatively confirm our hypothesis that a bifurcation is present in the myogenic differentiation trajectory, we simulated phase points in the RNA velocity vector field. Phase point analysis is a classical technique from dynamical systems that allows for quantitative descriptions of a vector field [82]. A “phase point” is initiated in the vector field, and its position is updated based on the vectors in the local neighborhood for many time steps. We initialized phase points on either side of the bifurcation point and updated their locations iteratively based on the RNA velocity of the nearest neighboring cells (Methods). We found phase points initiated after the bifurcation point (at a coordinate with greater pseudotime value) progress forward toward the differentiated attractor, while phase points initiated before the bifurcation point move backward toward the primitive *Pax7* high state (Fig. 5D). These results support the existence of a true bifurcation point that represents an energy barrier between maintaining a progenitor state (*Pax7* state) and committing to differentiation (*Tnnc1* state).

#### 2.6.1 Differentiation is delayed at the energy barrier in aged myogenic cells

Our analysis thus far provides a geographic understanding of the dynamics of myogenic differentiation as inferred by RNA velocity. However, it is difficult to identify differences in the young and aged velocity fields by purely qualitative analysis. To determine if young and aged cells progress differently through the differentiation trajectory, we projected the RNA velocity vectors into pseudotime along the differentiation trajectory and computed a ΔPseudotime coordinate for each cell (Methods)

Comparing the ΔPseudotime distribution across ages, we found that young cells had a broader distribution (Levene’s test, *p* < 0.001, Fig. 5E). To determine if there were differences in ΔPseudotime along the differentiation trajectory, we computed a rolling mean of ΔPseudotime values across pseudotime. Aged cells exhibited lower ΔPseudotime values near the bifurcation point, suggesting that it represents an energy barrier that aged cells are less able to overcome (Fig. 5F). This difference is significant considering all observed cells (Methods, nested linear models, log-likelihood ratio test, *p* < 0.001).

After the bifurcation point, aged cells exhibited only slightly lower ΔPseudotime values than young cells, but these may also create a meaningful difference in the pseudotime distribution (Fig. 3B). To test this latter hypothesis, we simulated phase points initiated after the bifurcation point and updated their locations based on either the young or aged velocity field. We found that phase points progress more slowly through the pseudotime trajectory in the aged velocity field (Fig. 5G). These results suggest that aged myogenic cells stall during the commitment decision and differentiate slower after committing.

To identify genes that may contribute to these observed differences in velocity between young and aged cells, we compared transcription, splicing, and degradation rates inferred for young and aged myogenic cells using a dynamical RNA velocity model [7] with a bootstrap resampling scheme (Methods). Genes with higher inferred transcription rates in young cells were enriched for the Hallmark Myogenesis gene set, while genes with different splicing and degradation rates across ages showed no enrichment (Fig. S24). This suggests that differences in differentiation rate across ages may be driven by differences in the transcription rates, rather than differences in splicing or degradation rates.

#### 2.6.2 Differential expression identifies molecular determinants of myogenic fate progression

The presence of a bifurcation point suggests that myogenic cells must decide to either progress forward in differentiation or reverse toward a primitive cell state. To determine which genes influence this decision, we calculated differential expression between cells moving “forward” and “backward” in the repulsor state. To identify cells moving forward and backward, we projected the RNA velocity vectors into pseudotime along the differentiation trajectory and computed a ΔPseudotime (Methods). To focus on cells with reliable direction estimates, we estimated confidence intervals for the RNA velocity vectors by parametric bootstrap sampling of UMIs for each cell. We then propagated these confidence intervals to the ΔPseudotime estimates (Fig. 6A, Methods). We only considered a cell to be moving forward if its 95% confidence interval for ΔPseudotime was greater than zero, and vice-versa for cells considered to be moving backward. There are 649 cells in the repulsor region with 391 cells moving forward (60%), 247 moving backward (38%), and 11 cells with confidence intervals crossing zero (2%).

**Figure 6:**
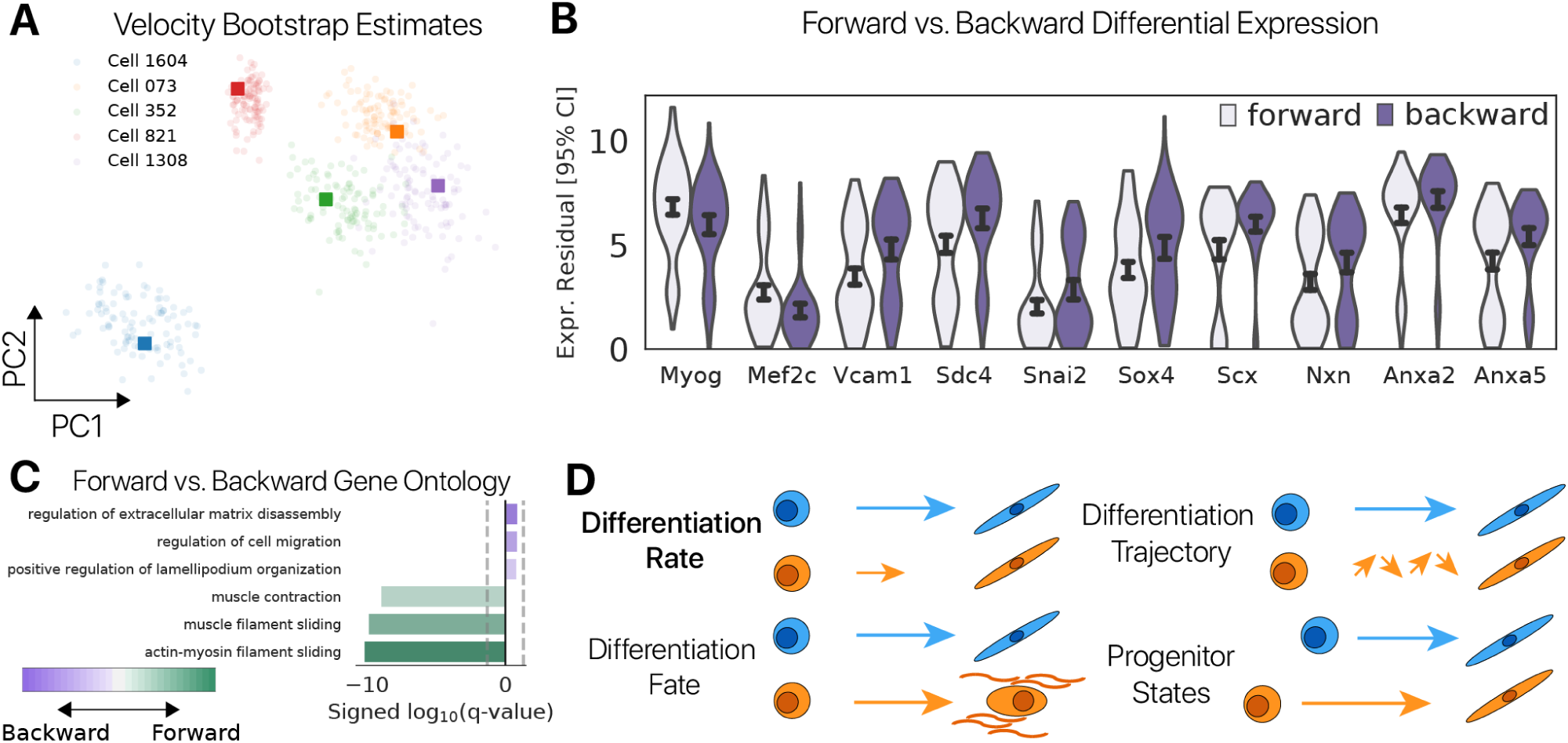
RNA velocity reveals molecular determinants of progression through the energy barrier. **(A)** PCA projection of RNA velocity vectors derived using our molecular resampling procedure. Colors indicate the cell of origin for each estimated vector. Circular points are estimates from a bootstrap sample, and square markers are velocity estimates computed with all observed reads. From the distribution of vectors, we derived RNA velocity confidence intervals. **(B)** Differentially expressed genes between cells moving forward and backward in the repulsor region. Values shown are residuals after regression of the pseudotime covariate from log(CPM + 1) counts and addition of minimum values to give a positive range. Myogenic transcription factors *Myog* and *Mef2c* mark cells progression forward, while primitive cell markers *Sdc4, Vcam1* and stress markers *Anxa2, Anxa5* mark cells moving backward. This suggests that relative levels of myogenic transcription factors, primitive cell markers, and stress signals may influence cellular decisions to move forward or backward through the differentiation program. **(C)** Gene Ontology enrichment analysis indicates that myogenic genes are downregulated in backward moving MuSCs relative to forward progressing MuSCs. **(D)** Schematic overview of conceptual models for the difference in differentiation outcomes between young and aged myogenic cells. Here, we found support for a difference in the rate of myogenic differentiation based on transcriptional distributions and dynamics (Differentiation Rates). We do not find evidence of a difference in cell fate with age or for differences in differentiation trajectories between young and aged cells. We likewise find that the differentiation state of young and aged progenitors is comparable. These results suggest that differences in the rate of differentiation at the transcriptional level underlie differences in the phenotypic outcomes.

Differential expression analysis of cells within the repulsor that are moving forward versus backward identified 129 genes (Methods). Cells moving forward were marked by myogenic transcription factors *Myog* and *Mef2c*, as well as muscle-specific cytoskeletal components such as *Actc1*. By contrast, cells moving backward had higher expression of primitive cell markers *Vcam1* and *Sdc4* [74, 18] and cell stress markers *Anxa2* and *Anxa5* (Fig. 6B). We also identified the transcription factors *Scx, Sox4*, and *Snai2* as markers of backward moving cells (Fig. 6B). *Snai2* is known to enforce a primitive identity during myogenic differentiation [76]. *Scx* and *Sox4* transcription factors have reported roles in myogenesis [52, 38] and may enforce a primitive myogenic identity at the bifurcation point. Genes upregulated in forward moving cells are enriched for myogenic gene sets, consistent with the idea that myogenic genes promoter commitment (Fig. 6C).

Forward and backward moving cells segregate in splicing phase space for velocity driver genes, but not for these genes (Fig. S23). This suggests the differentially expressed genes indirectly influence forward and backward motion, rather than directly driving the estimated ΔPseudotime. This is consistent with known biological roles for some differentially expressed genes. For example, *Snai2* is a known suppressor of *Myod1* [76], which activates some of the velocity driver genes we observe (e.g. *Dmd, Ttn, Macf1* either directly or through activation of *Myog*. Together, these results suggest that the relative levels of myogenic transcription factors and primitive cell signaling molecules dictate the direction of motion along the myogenic differentiation trajectory.

## 3 Discussion

Skeletal muscle is known to experience age-related decline in mass and regenerative potential, in part due to cell intrinsic defects in muscle stem cells. However, it remains unknown how age-related changes manifest across the differentiation trajectories of muscle-resident progenitors. Although, single cell expression studies have reported cell identity specific age-related changes in homeostatic tissues [2, 44, 83], it is difficult to determine which of these changes is associated with functional decline. Observing age-related changes during a functional challenge such as differentiation may reveal which differences are more closely associated with functional defects. Here, we used single cell RNA-seq to measure age-related changes across the differentiation trajectories of both muscle stem cells and muscle-resident fibro/adipose progenitors to investigate this possibility.

Our analyses revealed that age-related changes in both progenitor populations are only partially inherited by differentiated daughter cells, while differentiation revealed new age-related changes in daughters that are unseen in the progenitor pool. We observed this result at the level of individual genes (Fig. 3, Fig. 4) as well as in gene expression programs (Fig. 3I). To compare the overall magnitude of aging between two populations, we introduce a maximum mean discrepancy (MMD) based method that allows for rigorous statistical testing [30]. Quantifying the magnitude of age-related change before and after differentiation, we found that differentiation alters the overall difference between young and aged cells in a progenitor population (Fig. S9A; Fig. S10C). These results demonstrate that functional challenges such as differentiation can manifest age-related changes that are otherwise latent in quiescent, unchallenged cells.

Although aged MuSCs produced fewer late-stage differentiated cells, the overall trajectory of myogenic differentiation was conserved between young and aged MuSCs. Pseudotime analysis [87, 32] allowed us to establish a continuous, quantitative notion of lineage progression in both young and aged MuSCs leveraging the entire transcriptome, as opposed to a discretized set of differentiation states defined by marker genes. From these pseudotime trajectories, we found that young and aged differentiation trajectories were no more distinct than trajectories computed for different animals of the same age. This result suggests that aged MuSCs do not alter their differentiation trajectory in a cell-intrinsic fashion, as described for aged hematopoietic stem cells [5] and previously proposed for myogenic stem cells [13].

We observed that aged MuSCs exhibit delayed myogenic differentiation (Fig. 3B), suggesting that aging reduces the ability of aged stem cells to produce differentiated daughters. These observations are consistent with previous reports that aged MuSCs exhibit delayed cell cycle entry and reduced regenerative potential [28, 19]. RNA velocity inference [47] allowed us to move beyond this initial observation and identify the location in the differentiation trajectory where this delay occurs.

Adopting a concept from dynamical systems [82], divergence maps of the RNA velocity field revealed an unstable repulsor state separating two stable attractors, one marked by stem cell marker *Pax7* and the other by myocyte marker *Tnnc1* (Fig. 5B). Phase point simulations in the RNA velocity field indicate that the repulsor represents a bifurcation point where cells decide to travel toward the *Pax7* attractor or the *Tnnc1* attractor (Fig. 5D). Enabled by our method to estimate RNA velocity confidence intervals, we identified transcription factors and cell signaling genes that mark cells moving forward and backward at the repulsor as candidate molecular determinants of this cellular decision. Inferring a change in pseudotime along the differentiation trajectory (ΔPseudotime) for each cell, we found that aged MuSCs struggle to surpass this barrier. These analyses reveal the utility of dynamical systems formalisms applied in the context of single cell gene expression analysis, taking us from a phenomenological observation of reduced differentiation to a specific point in the trajectory where aged MuSCs are delayed. Our results support a conceptual model where myogenic differentiation outcomes change with age due to changes in the differentiation rate, rather than due to changes in cell fate or commitment trajectories (Fig. 6D).

Collectively, our results indicate that aging alters not only the baseline transcriptional state of progenitor cells, but also the transcriptional states that manifest throughout differentiation. Importantly, our experiments demonstrate that only a fraction of age-related changes in progenitors persist through differentiation, whereas differentiation simultaneously reveals age-related changes unseen in progenitors. Dedifferentiation to a pluripotent state has been reported to reverse many hallmarks of aging [48, 57, 89, 59, 71, 54], indicating that cell identity changes can mask age-related change in an extreme context. Our results further demonstrate that age-related changes can be modulated during the physiologically relevant cell identity changes in differentiation, suggesting that previous results in induced pluripotent systems are not a special case. Future work may explore a broader set of cell identity manipulations to reveal the causal relationships that underlie the plasticity of some age-related changes, and the perdurance of others.

## 4 Methods

### 4.1 Animals

All mice used were male, C57Bl/6 animals obtained from the Jackson Laboratories. Aged mice were received at Calico Life Sciences at 3 months of age and aged in-house until experimentation at 20 months of age. Young animals were received at Calico at two months of age and maintained in-house for one month to acclimate prior to experimentation at 3 months of age.

### 4.2 Cell Isolation

Prior to cell isolation, mice were perfused with 15 mL PBS using an automated pump to control flow rate (Harvard Apparatus). We isolated cells from muscle tissue as previously described [43, 72]. Briefly, we removed limb muscle tissue and minced using surgical scissors into pieces roughly 1 mm wide and 2-4 mm long. We incubated minced tissue in 15 mL, 2 mg/mL collagenase type II (Gibco) in DMEM for 60 minutes at 37^*o*^C with inversion of the tubes to mix every 5 minutes. We terminated this first digestion by dilution in wash buffer (F10 media, 10% horse serum) and followed by centrifugation (400 x *g*, 5 minutes) and a second wash. We incubated cells in a second digestion buffer containing 0.67 mg/mL dispase II (Gibco) and 0.13 mg/mL collagenase II in wash buffer for 30 minutes at 37^*o*^C with mixing every 5 minutes.

Post-digestion, we vortexed cells on high for 30 seconds to dislodge cells from fibers, centrifuged, and washed in wash buffer. We filtered the resulting suspension through a 40 *µ*m cell strainer and washed with wash buffer. After centrifugation and resuspension in wash buffer, we again filtered the cell suspension through a 40 *µ*m cell strainer into a 5 mL FACS tube and performed another wash. We then added FACS antibodies at the appropriate dilution and stained for 45 minutes on ice in the dark. Antibodies were washed out with 2 mL wash buffer at the end of the staining period. Prior to FACS, we also add propidium iodide (PI, dead cell marker) and Calcein Blue AM (Thermo-Fischer, live cell marker) following the manufacturer’s protocol.

### 4.3 Fluorescence activated cell sorting

To stain for muscle stem cells, we labeled cells with antibodies against CD31, CD34, VCAM1, Ly6a (Sca1), and *α*7-integrin. To positively mark fibro/adipose progenitors, we labeled cells with an antibody for PDGFR*α*. All cells were stained with the negative viability marker propidium iodide (PI) and positive viability marker Calcein Blue AM (Thermo Fischer). Antibodies are described in Table S1.

For the initial timepoint with cells from the whole muscle mononuclear cells suspension, MuSCs were enriched by gating on CD31^-^/CD45^-^/Ly6a^-^/VCAM1^+^/*α*7-integrin^+^ and sorting out 10,000 cells. FAPs were enriched by gating on CD31^-^/CD45^-^/Ly6a^+^/PDGFR*α*^+^ and sorting 10,000 cells. Blood cells were enriched by sorting 10,000 cells from the CD31^+^/CD45^+^ population. A bulk population of live cells using the viability markers was sorted after these populations have been collected. For cells at the differentiated timepoint, we sorted for live cells.

### 4.4 Cell Culture

Sorted MuSCs and FAPs from the initial “quiescent” timepoint were seeded into separate wells of a 96-well place (coated with sarcoma-derived ECM, Sigma Aldrich). MuSCs were seeded with 3,000 cells per well with 6 wells per animal, while FAPs were seeded with 5,000 cells per well with 6 wells per animal. Both populations were maintained in DMEM with 10% horse serum and penicillin/streptomycin (Gibco). This media has been commonly used to induce differentiation in both populations over a 2-4 day period in the literature [16, 39]. Cells were maintained in a 5% CO_2_, 37^*o*^C incubator for 60 hours prior to dissociation. After initial seeding and prior to dissociation, cells were imaged by brightfield microscopy to capture morphological distinctions. We dissociated differentiated cells from the culture dish by incubation with 0.25X Trypsin EDTA (Gibco) for 2 minutes, followed by pipette mixing until cells were dislodged. We mixed dissociated cells into a single cell suspension for each animal prior to live cell sorting and single cell RNA-sequencing.

### 4.5 Single cell RNA-sequencing

For the initial “quiescent” timepoint, we mixed 5,000 sorted MuSCs, 5,000 sorted FAPs, 5,000 sorted blood cells, and 15,000 cells from the bulk suspension to create a cell suspension enriched for MuSCs, FAPs, and blood cells. We immediately transferred 10,000 cells from each of these suspensions to a 10X Chromium instrument for emulsion using a 3’ Single Cell Transcriptomics v3 kit (10X Genomics). For the second “differentiated” timepoint, we loaded 10,000 live cells per well. A single 10X Chromium well was used for each animal at each timepoint. Following emulsion, library preparation was prepared following the 10X Genomics Single Cell Transcriptomics v3 protocol. All samples were evaluated for quality using an Agilent Bioanalyzer prior to sequencing.

### 4.6 Read Alignment and Gene Expression Quantification

We aligned reads to the `mm10` reference genome from Ensembl. We used genome release 97, GRCm38.p6 with slight modifications to the annotations as previously described [44]. We replaced annotations for lincRNAs *Gm42418* and *AY036118* with a single contiguous gene annotation for the rRNA element *Rn45s* on chromosome 17 beginning at 39,842,000 bp and ending at 39,850,000 bp. We prepared a cellranger reference using `cellranger mkref`. We performed alignment, cell barcode demultiplexing, UMI demultiplexing, and cell barcode filtering using cellranger 3.0.2 from 10X Genomics [96]. cellranger makes use of the STAR sequence aligner [22]. This analysis yields a Cells × Genes matrix, where the *i, j*-th element is the number of UMIs detected in cell *i* mapping to gene *j*. We removed pseudogenes and single exon lincRNAs from this count matrix prior to subsequent analysis.

### 4.7 Quality Control

We removed cell barcodes with a high proportion of reads mapping to the mitochondiral genome > 20% as putative dead cells [36]. Genes detected in fewer than 10 cells were removed from downstream analysis.

### 4.8 Doublet Removal

We identify doublets using the `solo` neural network classification scheme [9]. First, we fit a single cell Variation Inference (scVI) [53] model to our raw counts matrix. We then create “synthetic doublets” by randomly sampling 2 cells at a time from our counts matrix and adding their expression profiles. We then embed these “synthetic doublets” in the latent space learned by the scVI model and train a neural network classifier to discriminate these synthetic doublets from real cells. This trained classifier is roughly 87% accurate on our data set. We subsequently classify all the real cell samples using this classifier and remove predicted doublets.

### 4.9 Normalization and Dimensionality Reduction

Raw UMI counts were normalized by the library size to yield Counts Per Million (CPM) and log transformed after adding a pseudocount of 1 (log(CPM+1)). Differential expression results were computed using these log-normalized counts. To construct an embedding, we normalized raw cell counts using a regularized negative binomial regression approach implemented in `sctransform`, yielding Pearson residuals [31]. We selected the 1,500 most highly variable genes (HVGs) in the Pearson residuals based on empirical variance within the gene expression vector. We constructed a nearest neighbor graph in this HVG space using the 15 nearest neighbors and performed a UMAP embedding to generate a 2D projection of the high-dimensional data with parameters `min_dist = 0.5` and `spread = 1.0` [56, 4].

### 4.10 Clustering and Cell Type Identification

We performed Leiden community detection [85] on the nearest neighbor graph to identify a cluster partition. We inferred cell types using a deep neural network classifier trained on data from the *Tabula Muris* [73]. Our classification approach is similar to our previous report, with slight modification [44]. The classifier used 4 fully-connected neural network layers, each followed by a rectified linear unit activation, batch normalization, and a dropout layer. The two hidden layers adopted a residual architecture as introduced for Residual Networks [35], each with *n* = 256 units. A dropout layer was also applied to each input with probability *p* = 0.3 prior to the first layer to simulate technical noise during mRNA capture and measurement and improve generalization. During training, we perform class balancing by random sampling to remove bias due to cell type proportions. Classes with fewer than 128 examples are oversampled, while classes with more examples are undersampled. Models were trained with the Adagrad optimizer [23].

We trained a classifier to predict cell types using “cell ontology class” annotations provided for `Limb Muscle` data in the *Tabula Muris*. We refined cell type predictions from this model by smoothing class labels to the modal class of a *k* = 15 neighborhood around each cell, where nearest neighbors were restricted to cells within the same cluster. This post-processing step allows neighborhood information to aid in cell type identification, and has been reported to improve cell type inference using other models [64]. Cell type annotations were validated manually using marker gene expression. Our experimental data also contains cell types not annotated in the *Tabula Muris*. We annotated these cell types manually by examining marker gene expression.

### 4.11 Differential Expression

Differential expression analysis for binary contrasts (e.g. young vs. aged) was performed using the Wilcoxon rank sums test on log(CPM + 1) normalized counts. The false discovery rate was controlled at *α* = 0.05 using the Benjamini-Hochberg procedure [6].

Common differentially expressed genes were identified by first quantifying the number of unique cell identities in which each gene was differentially expressed in the same direction with age. Genes that were expressed in *k* >= 3 cell identities were considered to be common differentially expressed genes.

Differential expression for continuous covariates (e.g. pseudotime) was performed using nested hurdle models. We constructed hurdle regression models using a combination of a logistic regression and Gaussian generalized linear model (GLM), as described for the MAST method [26]. Briefly, for each gene expression vector *y* of log(CPM+ 1) counts, we define a binary indicator vector *z* = 𝟙_*y*>0_. We fit full and reduced logistic regression models to *z* for each gene, where reduced models are nested within full models. We then fit full and reduced Gaussian GLMs to *y* only where *z* ≡ 1. The logistic regressions therefore describe the probability a given gene will be expressed using provided covariates, while the Gaussian GLM models describe the expected expression level conditioned on the gene being expressed. As in MAST, we include the “cell detection rate” (CDR, fraction of genes detected in each cell) as a nuisance covariate in our regressions [26].

For significance testing, we perform a log-likelihood ratio test between our full model and reduced model, using the *χ*^2^ distribution to determine *p*-values as in Monocle 2 [65]. We performed multiple hypothesis test correction using the Benjamini-Hochberg procedure as above. This scheme allows us to test the influence of an additional covariate and its interaction terms on the expression of a given gene. To identify genes significantly influenced by pseudotime, we fit a full model *y* ∼ Pseudotime + Age + Age:Pseudotime + CDR with a corresponding reduced model *y* ∼ Age + CDR. To identify genes with a significant age:pseudotime interaction, we fit full models *y* ∼ Pseudotime + Age + Age:Pseudotime + CDR with a corresponding reduced model *y* ∼ Pseudotime + Age + CDR. To identify genes significantly influenced by age after controlling for pseudotime distribution, we fit full models *y* ∼ Pseudotime + Age + Age:Pseudotime + CDR with corresponding reduced models *y* ∼ Pseudotime + CDR.

### 4.12 Local Fold-Change Estimation

We estimated the fold-change for features of interest **z** (e.g. gene, gene expression program activity) with respect to cell state using the nearest neighbor graph. We first defined a binary contrast variable **y** ⟶ {0, 1} for computing fold changes. We then constructed a *k*-nearest neighbor graph in principal component space using only cells specific to each level of the contrast variable, yielding two graphs *G*_0_ and *G*_1_. For each cell *i* in our observations, we identified the *k* nearest neighbors in each of the two graphs and computed the fold-change in the feature of interest **z**. We only compute a fold-change for a given cell if the number of cells expressing a minimum value of the feature of interest *t* is larger than a threshold number of cells *m*. If this criterion is not met, we assign a fold-change of 0. for visualization purposes.

For analyses presented here, we used *k* = 100 nearest neighbors. We set the threshold of expression *t* = 0 for individual genes with *m* = 30 as a minimum number of cells passing the threshold. For gene expression programs, we set the threshold *t* = Otsu(**z**) where Otsu is the Otsu thresholding function [60] and *m* = 50 as the minimum number of cells. We increase the minimum number of cells, as the a gene expression program score is less prone to a failure of detection than a single gene.

### 4.13 Non-negative Matrix Factorization

We perform non-negative matrix factorization (NMF) on a matrix of quiescent and differentiated myogenic cells and FAPs containing the 2,000 top highly variable genes (selected using over-dispersion) [91]. For optimization, we use random initialization and multiplicative updates [49]. We used an NMF embedding with *k* = 20 dimensions to mitigate the “curse of dimensionality” [34] (58.3% of variance explained, Fig. S15C).

To interpret the gene expression programs captured in each latent variable, we thresholded the gene loadings using Otsu thresholding [60] to assign a set of genes to each latent variable. From these genes, we identified enriched Gene Ontology terms from the Biological Process gene set library. We derived a semantic name for each latent variable based on these gene set enrichment results.

### 4.14 Aging Trajectories

We computed aging trajectories as the distance between the centroids of young cells *µ*_*Y*_ and aged cells *µ*_*A*_ in the NMF embedding, 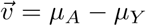. To analyze aging trajectories in a continuous fashion along a pseudotime trajectory, we ordered cells according to their pseudotime coordinates and computed aging trajectories within a sliding window of *k* = 500 cells with a stride *s* = 10.

### 4.15 Estimating Aging Magnitude by Maximum Mean Discrepancy

We estimate the magnitude of aging in a cell population using the maximum mean discrepancy (MMD) metric. Briefly, MMD attempts to determine the difference between two distributions *p*(*x*) and *q*(*y*) given only samples from the distributions, *x* ∼ *p*(*x*); *y* ∼ *p*(*y*). We can estimate the difference between these two distributions by computing the difference in the expectations of the samples after passing through some function *f*. This mean discrepancy – the difference between the expectations – reduces to a simple difference in means in the case where *f* is an identity function. If we search over a sufficiently flexible family of functions *f*, we may expect that the maximum diffference in expectations will only be 0 if *p* ≡ *q*, whereas the maximum mean discrepancy across this family of functions would be larger than 0 if *p* ≠ *q*. This can be shown in the case where *f* is a kernel metric, commonly implemented as a radial basis function (RBF) kernel [30]. We leverage this case and compute the MMD as

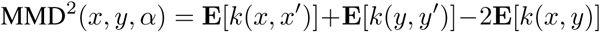

where *x, x*′ ∼ *p*(*x*), *y, y*′ ∼ *q*(*y*), and *k*(*x, y*) = exp(−*α‖x − y*‖^2^). We set *α* to the median pairwise distance between samples and computed MMD on random samples of 500 cells per age across several iterations, which we refer to as a heterochronic comparison.

We also computed the MMD on different random samples from the same age to generate an empirical null distribution (e.g. comparing random samples of young cells to young cells), which we refer to as isochronic comparisons. We employ the kernel two sample test to assess the statistical significance of the MMD [30]. We performed our MMD comparisons using log CPM + 1 normalized transcript counts, omitting genes expressed in < 40 cells (≈ 0.16% of measured cells) across the entire data set to reduce dimensionality.

### 4.16 Cell Age Classification and Latent Space Exploration

Deep neural network (DNN) classifiers were trained to predict cell age. Each model used a single residual block [35] with 128 hidden units. We optimized a weighted cross-entropy loss with Adagrad, weighting minority classes more heavily (weight *u*_*k*_ = 1*/p*(*K* = *k*) where *p*(*K* = *k*) is the proportion of input cells in class *k*). For dynamic data augmentation, we used MixUp [95] to interpolate between observed data points, sampling the MixUp weight parameter *γ* from a Beta distribution with shape parameters *α* = 0.2. We also dynamically augmented input vectors by randomly setting 30% of the input gene counts to 0 at each epoch. We applied a weight decay of 0.0001 during training and used batch sizes *M* = 128. For pericytes and Schwann cells, we increased regularization by setting weight decay to 0.01, MixUp *α* = 0.5, and dynamically setting 60% of input gene counts to 0 to prevent overfitting on the small sample sizes.

Models were trained independently for each cell type and time point to discriminate young and aged cells. Each model was trained using 5-fold cross-validation, with separate training, validation, and testing sets. The validation set was used for model selection by early stopping, while the testing set was held-out and unseen, used only for evaluation of the trained model. We report age classification accuracies as the mean testing accuracy across cross-validation folds.

DNN age classifiers learn an embedding where young and aged cells are easily separable with a linear classifier (the final layer of the DNN model). To explore these embeddings, we predict the age of all cells in a population and extract the neuron activations from the penultimate layer. For presentation, we project the latent space into 2-dimensions using UMAP.

We identified salient genes that drive classification decisions using guided backpropogation [78]. Briefly, given a trained classification model *f*_*θ*_ we computed the gradient on a target class score *f*_*θ*_(*x*)_*k*_ with respect to a set of input cells *x* in the target class *k*. At each rectified linear unit (ReLU) in the model *f*, we rectify the gradient during backpropogation by passing the local gradient *g* through max(0, *g*). We present features with the highest mean saliency across the set of input cells. For each class, we use a random sample of 50 cells from the target class *k* ∈ {Young, Aged}.

### 4.17 Identifying Cycling and Apoptotic Cells

We identified cycling cells using a standard gene set scoring method [84]. Briefly, a set of genes associated with S-phase and G2/M-phase cells was derived from single cell RNA-sequencing in 293T and 3T3 cell lines. For each cell, the sum of the expression of these phase-associated genes were computed. As a null distribution, a set of genes were selected from the set difference of the observed genes and the phase-associated genes. These null genes were selected by binning the sample genes into 24 equally sized bins by mean expression, then randomly sampling 100 null genes per bin per phase-associated gene with replacement. The difference between the mean of the phase-associated genes and the selected null genes was considered the module score. From these scores, we identified a cluster of myogenic cells enriched for cell cycle gene expression as cycling cells.

To identify apoptotic cells, we scored the activity of the “Hallmark Apoptosis” gene set from MSigDB and the “apoptotic process” gene set from the Gene Ontology database using the AUCell method [1]. We performed Leiden community detection to isolate a cluster of myogenic cells with higher activity in these two apoptotic gene sets and removed these cells from pseudotime analysis.

### 4.18 Pseudotemporal Trajectory Fitting

We inferred pseudotime trajectories using the diffusion pseudotime method [33]. Diffusion pseudotime requires the selection of a root cell to use as the diffusion source. We selected root cells in the myogenic differentiation trajectory by clustering cells with Leiden community detection and selecting a subset of cells with high *Pax7* expression as potential root cells. *Pax7* is a known marker of the myogenic stem cell state [14]. A root cell was selected randomly within this population. We fit a common pseudotime trajectory to aged and young cells to compare the distribution of cells along the pseudotime trajectory.

To compare cells from early and late periods in the myogenic pseudotime trajectory, we ordered cells by their pseudotime coordinates and binned them into 3 equal width bins. We considered the first bin to be “early” in differentiation and the final bin to be “late.”

We selected a root cell in the fibro/adipose progenitor population based on high expression of *Cd34*, a mesenchymal progenitor cell marker. We fit a single diffusion pseudotime trajectory for all cells in the differentiated FAP population, then identified distinct adipocyte and myofibroblast differentiation trajectories with PAGA [90]. We performed Leiden community detection to identify coarse cell states, then used PAGA to build a graph between these states. We defined paths through this coarse state graph corresponding to adipocyte and myofibroblast fates, confirmed using marker genes. For each fate, we defined a pseudotime covariate corresponding to the unique differentiation trajectory of that fate by taking only cells along the corresponding PAGA path and normalizing pseudotime values among these cells [0, 1]. We projected the FAP differentiation trajectories into a 2-dimensional embedding using ForceAtlas2 [37] with the positions of coarse cell states in the PAGA graph as initial positions.

### 4.19 Pseudotemporal Trajectory Gene Kinetic Curves

We fit quartic splines using `SciPy UnivariateSpline` to model the expression of each gene’s log-normalized counts *y* relative to the pseudotime coordinates *x*. To identify genes with similar kinetics in the myogenic program, we fit a common set of splines across cells of both ages and inferred 100 points linearly spaced along the pseudotime trajectory for each gene. We clustered these spline-derived gene expression curves using hierarchical clustering to identify Gene Modules with similar kinetic behavior. We optimized the number of clusters *k* based on the Silhouette index.

We subsequently fit separate splines as above to young cells and aged cells for visualization. To estimate variation in these spline fits, we performed bootstrap resampling and fit splines to 90% of the cells of each age across 100 iterations. This procedure was repeated for the adipocyte and myofibroblast pseudotime trajectories in the FAP population.

### 4.20 Pseudotemporal Trajectory Distance Comparison

We use a dynamic time warping (DTW) distance [41] to compare pseudotime trajectories fit independently to aged and young myogenic cell populations. This method is similar to a pseudotime alignment method described previously [15]. For each age, we randomly sample a subset of *n* = 250 cells to allow for bootstrapping and to control for any effects of differing sample sizes between ages. Within this sample, we choose a root cell from the population described above and fit a diffusion pseudotime trajectory. Some pseudotime fits leads to infinite pseudotime values for a small number of cells (< 20) that are distant from the rest of the nearest neighbor graph. We remove these cells from the graph and refit the pseudotime trajectory to reduce effects from these outlier cells.

For each gene, we then fit a smooth spline *S* (using `scikit-learn UnivariateSpline`) after centering the gene expression to *µ* = 0 and scaling variance to *σ*^2^ = 1. From this smooth spline, we interpolate 100 points evenly spaced along the pseudotime axis from [0, 1]. Once smooth splines are computed for all genes for each age, we compute the DTW distance between splines for each gene between the two ages. We compute trajectory distance as the sum of gene-wise DTW distances.

### 4.21 RNA Velocity Estimation

We count the number of reads mapping to spliced and unspliced transcripts using `velocyto` [47]. Prior to fitting the RNA velocity model, we subset the Cells × Genes matrix to genes that have both a spliced and unspliced count in at least 20 cells. We then further select the 4000 most highly variable genes based on over-dispersion [91]. We fit the first moments of the RNA velocity model using *k* = 100 nearest neighbors after constructing a nearest neighbor graph using the first 30 principal components of the subset genes. Subsequently, we fit the deterministic RNA velocity model using the `scvelo` implementation [47, 7]. We computed velocity vectors in PCA embeddings by projection with the principal components weight matrix. For visualization, we embed velocity vectors in ForceAtlas2 graphical layouts using a *k*-nearest neighbors regressor (*k* = 30) trained to predict ForceAtlas2 layout coordinates from the first 10 principal component scores.

### 4.22 Gene Contributions to RNA Velocity Vectors

We estimated the importance of individual genes to the velocity vectors in the PCA embedding using the contributions of individual gene velocities to the principal component coordinates. We used the first principal component in the myogenic cell embedding to estimate contributions to “forward” and “backward” velocity, as this component is well-aligned with myogenic pseudotime.

We computed the elementwise product of gene-specific velocities in each cell **v**_*i*_ with the loadings of a principal component **W**_*j*_ to obtain a vector of “contribution scores” for each gene in each cell **s**_*i*_.

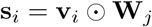

where ⊙ is elementwise multiplication. We computed the mean contribution scores across cells for each gene *g* as

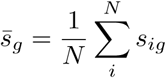

and then rank ordered the mean contribution scores for all genes. We selected the *k* = 100 genes with the most positive scores, and the *k* = 100 genes with the most negative scores as “forward” and “backward” driver genes respectively. For myogenic cells, we identified “forward” driver genes using cells from the latter portions of the pseudotime trajectory (Pseudotime > 0.4) and “backward” driver genes using cells from the early portions of the pseudotime trajectory (Pseudotime < 0.3).

### 4.23 RNA Velocity Divergence

We estimated the divergence of RNA velocity fields by coarse-graining the first two principal components of the cell embedding into *n* = 30 equally sized bins for each dimension, yielding a 30 × 30 grid of PC coordinates *X*^30×30×*D*^ where *D* = 2. In each bin *X*_*i,j*_, we computed the mean RNA velocity vector 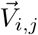 using a Gaussian kernel *𝒩* (*µ* = 0, *σ* = 0.5) to weight the contribution of the *k* = 100 nearest neighbor cells. We shrank the velocity vector to 0 for bins with few nearby cells (below the 5^th^ percentile of total probability mass, as computed using the Gaussian kernel). Our implementation borrows from a similar scheme used by `scvelo` [7]. This is a conservative post-processing step to avoid making divergence claims for portions of the latent space where we have few measurements.

We computed the divergence of the resulting vector field 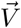 as:

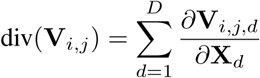

where *d* ∈ *D* are coordinate dimensions of the embedding.

To determine if the divergence of each bin in the coarse-grained embedding is non-random, we performef the above operations for 100 random samples of cells, each containing 80% of the total population. For each bin, we computed a 95% confidence interval for the divergence value using the bootstrap samples. We shrank the divergence value for a given bin *d*_*i,j*_ to 0 if this confidence interval contains 0.

### 4.24 RNA Velocity Confidence Intervals

We generated confidence intervals for RNA velocity vectors using a parametric bootstrapping procedure. For each cell, we fit a multinomial distribution to the mRNA abundance profile Multi(‖**x**_**i**_‖, **x**_**i**_*/*‖**x**_**i**_‖), where *x*_*ij*_ are the total counts for gene *j* in cell *i*. From this distribution, we drew bootstrap samples

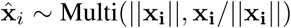

to reflect stochasticity in the random sampling process of RNA-seq, which captures a relative rather than absolute abundance profile.

For each gene in each cell, we fit a second multinomial to reflect the proportion of read counts mapping to that gene that originate from spliced, unspliced, or ambiguous transcripts, which we denote *x*_*ijs*_, *x*_*iju*_, *x*_*ija*_. We adopt the spliced, unspliced, and ambiguous definitions of La Manno *et. al*. [47]. From a second multinomial, we drew a sample of spliced, unspliced, and ambiguous reads given a sampled quantity of total reads

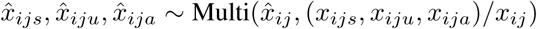

After performing this sampling procedure for all cells in an experiment, we fit RNA velocity estimates to the sampled reads as above. We repeated this procedure 100 times to estimate confidence intervals on RNA velocity vectors.

### 4.25 Dynamical Systems Simulation of Kinetics

To determine how a cell might progress through transcriptional space using RNA velocity estimates, we simulated the progression of a phase point in the RNA velocity vector field. We selected a set of initial starting locations by sampling the location of cells in the PCA embedding. For myogenic phase point simulations, we sampled cell locations that were either (1) within a region we identified as a “repulsor” or (2) beyond this region, where we expect cells to progress forward in pseudotime.

For each phase point, we selected a starting location *x*_*t*=0_ at random from this set. We determined the new location of the phase point at each step of the evolution process as:

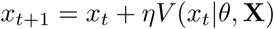

where *V* (*x*_*t*_|*θ*, **X**) is a function that estimates the RNA velocity at a location *x*_*t*_ in the embedding space given parameters *θ* = (*k, s*) and the observed embedding positions and velocity vectors **X**. We implemented *V* as a stochastic function that samples an RNA velocity vector using information from the observed *k*-nearest neighbors of the point *x*_*t*_. We estimated a weighted mean *µ*\* and weighted covariance matrix Σ* from the RNA velocity vectors of these *k*-nearest neighbors. We determined weights for each nearest neighbor using a Gaussian kernel *𝒩* _weight_(*x*_*t*_, *s***I**), where *s* is a scale parameter. We set *s* to the mean distance between nearest neighbors in the whole observed cell population.

We compute the weighted mean and covariance as:

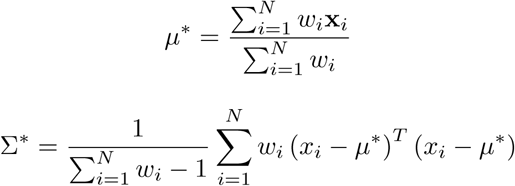

Using the weighted mean and covariance matrix, we instantiated a multivariate normal distribution 𝒩_sample_(*µ*\*,Σ*). We sampled a stochastic estimate of RNA velocity 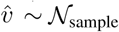 from such a distribution at each update step. We set the step size coefficient *η* = 0.01 for all simulations. We perform simulations for 1,000 phase points in each condition.

### 4.26 Change in Pseudotime Estimation

To estimate the “change in pseudotime” (ΔPseudotime) for a cell *i* with coordinates in the PCA embedding 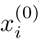, we estimate the cell’s future location in the embedding space

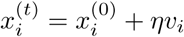

where *v*_*i*_ is the RNA velocity estimate for cell *i* in the embedding and *η* is a step size parameter we set to *η* = 0.01 for all predictions.

We train a *k*-nearest neighbors regression model 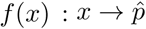 to estimate pseudotime coordinates 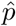 from principal component coordinates *x*. Using this model, we compute the change in pseudotime for cell *i* as:

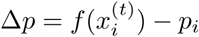

where *p*_*i*_ is the measured pseudotime coordinate in cell *i*. We computed confidence intervals on these Δ*p* coordinates by computing Δ*p* for each of 100 parametric bootstrap estimates of RNA velocity for each cell, described above.

### 4.27 Change in Pseudotime Significance Testing

To evaluate the significance of changes in ΔPseudotime predictions, we fit nested linear models and performed a log-likelihood ratio test. We fit a full regression of the form ΔPseudotime ∼ Pseudotime + Age + Age:Pseudotime + CDR with a reduced model of the form ΔPseudotime ∼ Pseudotime + CDR to evaluate the impact of age on ΔPseudotime in myogenic cells.

### 4.28 Differential Expression Based on Change in Pseudotime Direction

To identify cells that are moving forward or backward in pseudotime, we predicted ΔPseudotime coordinates for each of 100 bootstrapped estimates of RNA velocity (see above) and built a 95% confidence interval from these predictions. We considered a cell to be moving forward if and only if the 95% confidence interval of ΔPseudotime was greater than 0, and likewise a cell was considered to be moving backward if and only if the 95% confidence interval of ΔPseudotime was less than 0. To identify markers of forward and backward progression, we performed differential expression tests on the forward and backward labels for cells in the repulsor region after using linear models to regress out the pseudotime coordinate. We cast all residuals from this regression procedure to positive values by addition of the minimum expression value for each gene to avoid undefined values in fold-change calculations.

### 4.29 Change in RNA Velocity Parameters with Age

We estimated RNA velocity transcription (*α*), splicing (*β*), and degradation (*γ*) rates for each gene using the dynamical RNA velocity model in `scvelo` [7]. We excluded genes with < 20 total UMI counts from estimation, and subset to the 4000 most highly variable genes based on dispersion before model fitting. We built the `scvelo` nearest neighbor graph using the first 30 principal components of the spliced counts matrix and first *k* = 100 nearest neighbors.

To compare the distribution of parameter values between young and aged cells, we fit separate models to cells of each age. In order to compare parameter values for individual genes, we fit dynamical models to bootstrap samples of cells from each age to estimate the variation in a given parameter value. We randomly sampled 80% of cells from each age and fit 100 separate dynamical models to these bootstrap samples. We used Gene Set Enrichment Analysis (GSEA) and the MSigDB Hallmark gene sets determine which biological processes had altered parameter values with age.

## 5 Data Availability

Our data have been submitted to the NCBI Gene Expression Omnibus under accession number GSE145256. We have additionally made our data and software tools available at myo.research.calicolabs.com.

## 6 Acknowledgements

The authors would like to thank Calico Physiology, including Freeman Lewis, Chunlian Zhang, and Ganesh Koulumam for their assistance with animal and cell isolation experiments. The authors also thank members of Calico Genomics, including Margaret Roy, Andrea Ireland, and Twaritha Vijay for assistance with single cell RNA-sequencing experiments. We also thank David Botstein and Cynthia Kenyon for helpful discussions.

## 7 Author Contributions

JCK performed experiments and analysis. DGH guided analysis and supervised research. DRK guided analysis and supervised research.

---

**Figure S1:**
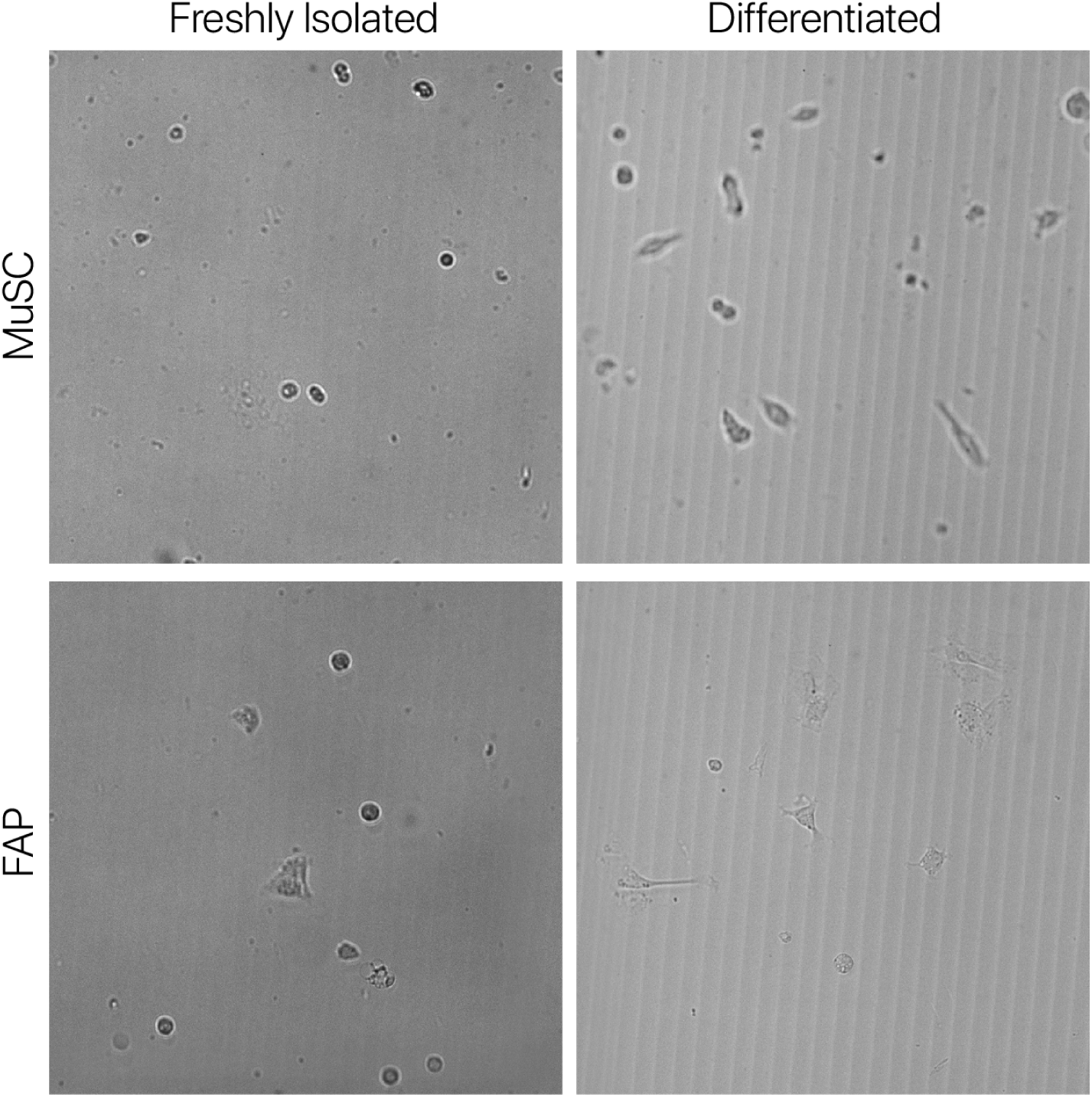
Representative images of freshly-isolated and differentiated MuSCs and FAPs *in vitro*.

**Figure S2:**
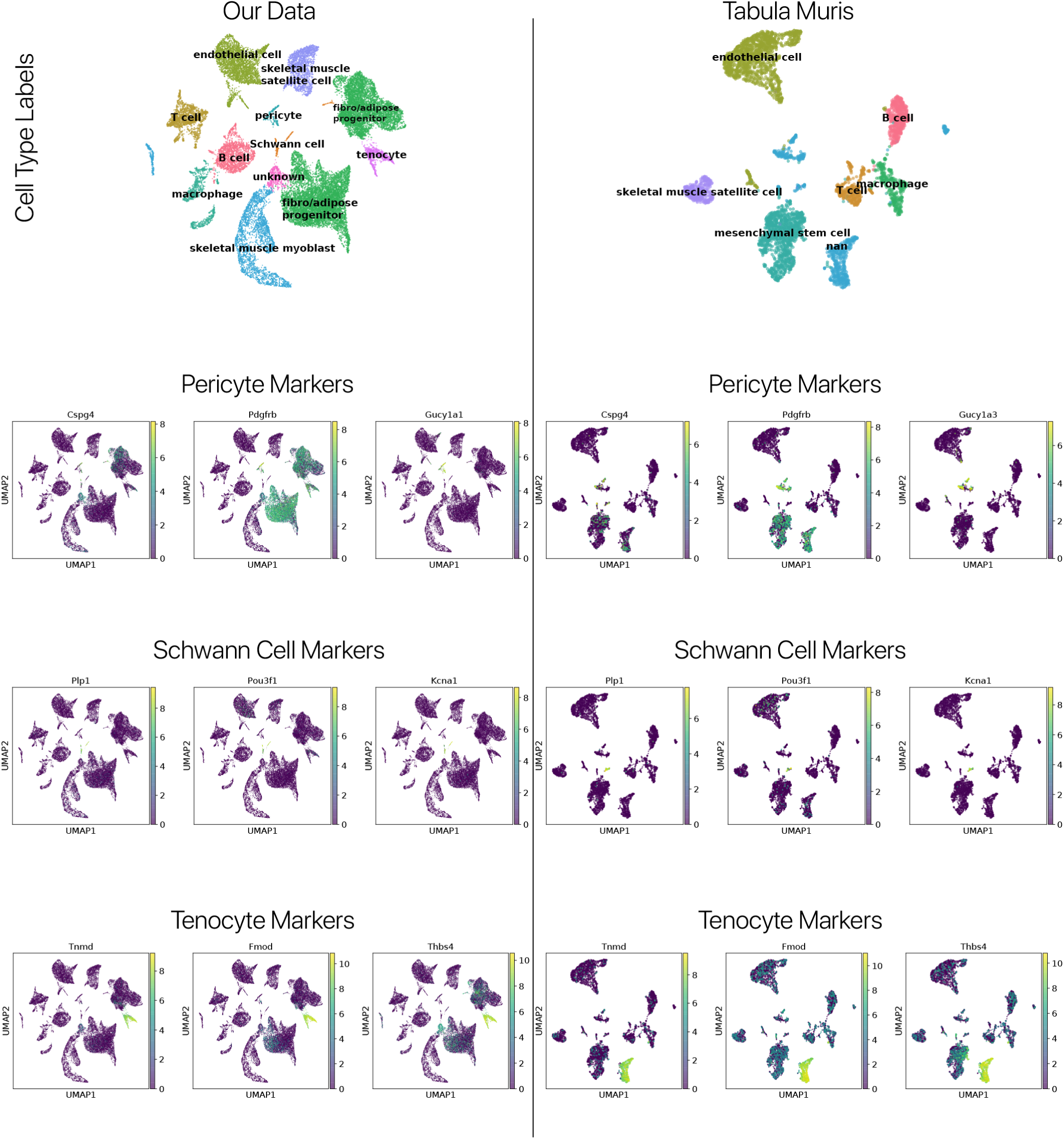
Identification of rare cell types based on marker genes in our data and the *Tabula Muris*. Cell type labels for our data and the *Tabula Muris* are shown. Note that annotations of “nan” in the *Tabula Muris* correspond to unannotated cells. Below, we show the expression of pericyte, Schwann cell, and tenocyte marker genes in both data sets. We find that rare cell types are present in both data sets, but provide novel annotations.

**Figure S3:**
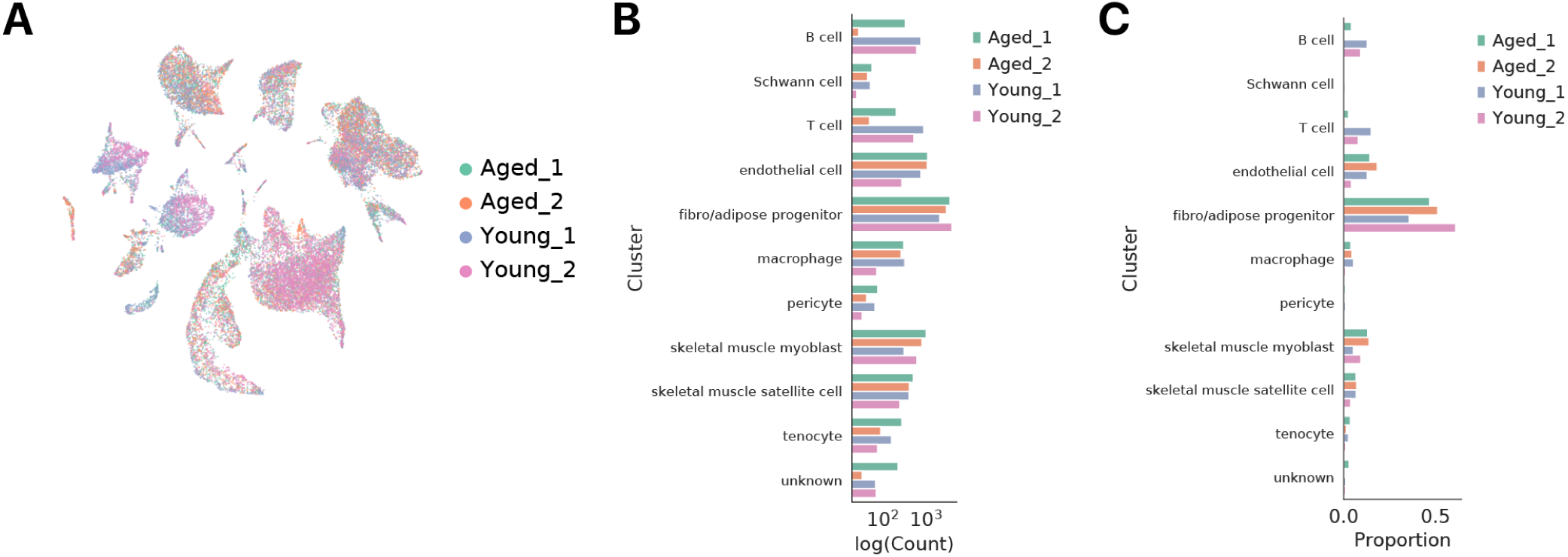
Each cell type is recovered in each animal. **(A)** Animal labels overlaid as colors on a UMAP projection of transcriptional space. Note that each cluster has cells from each animal. **(B)** Cell counts of each type for each animal. Note that we recover cells of each type in each animal. **(C)** Cell counts in **(B)** converted to proportions.

**Figure S4:**
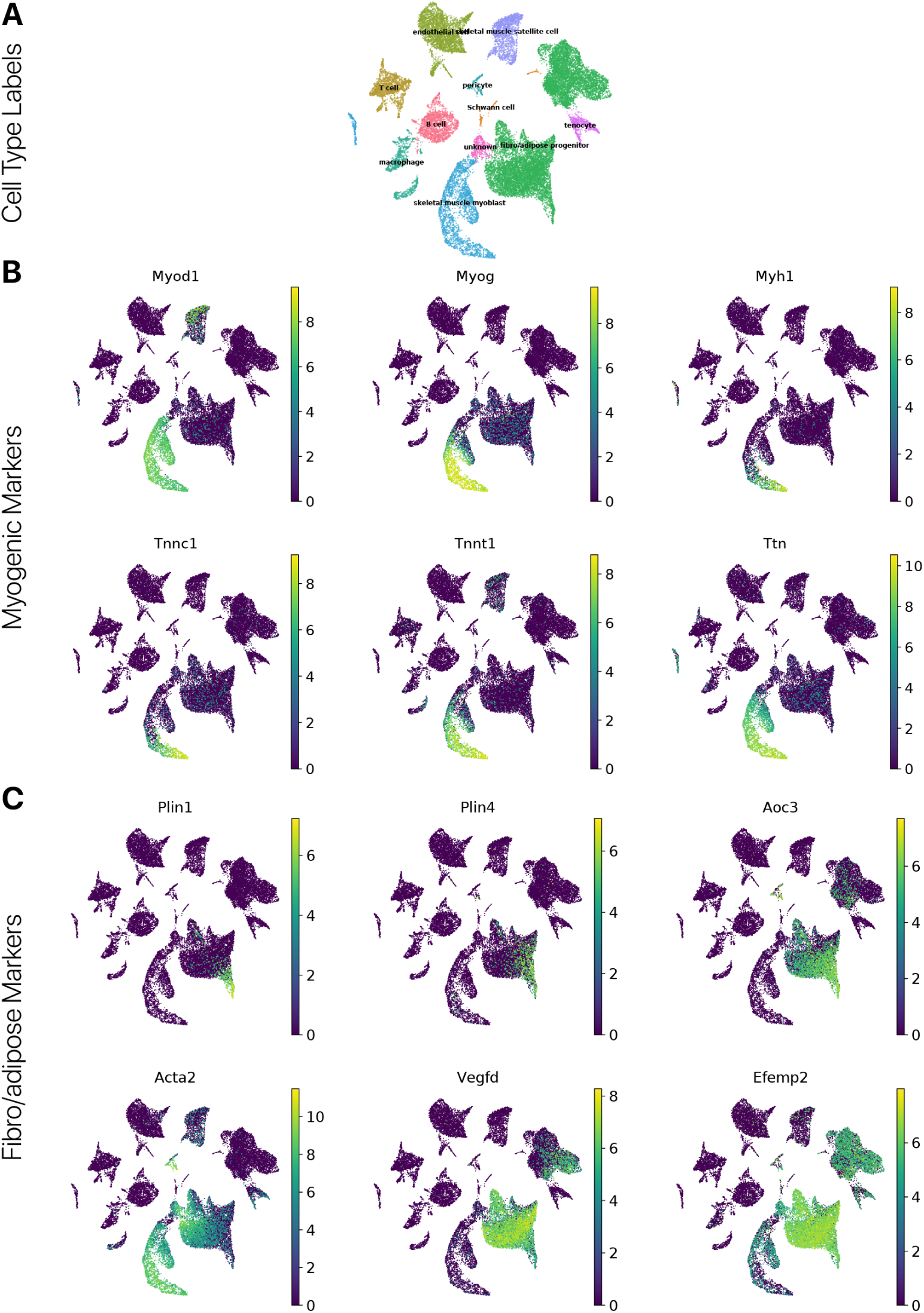
Differentiation marker genes confirm the efficacy of *in vitro* differentiation. **(A)** Cell type labels overlaid on a UMAP projection of transcriptional space. **(B)** Myogenic differentiation markers, enriched in myogenic cells from the differentiated time point. **(C)** Fibro/adipose differentiation markers, enriched in FAPs from the differentiated time point.

**Figure S5:**
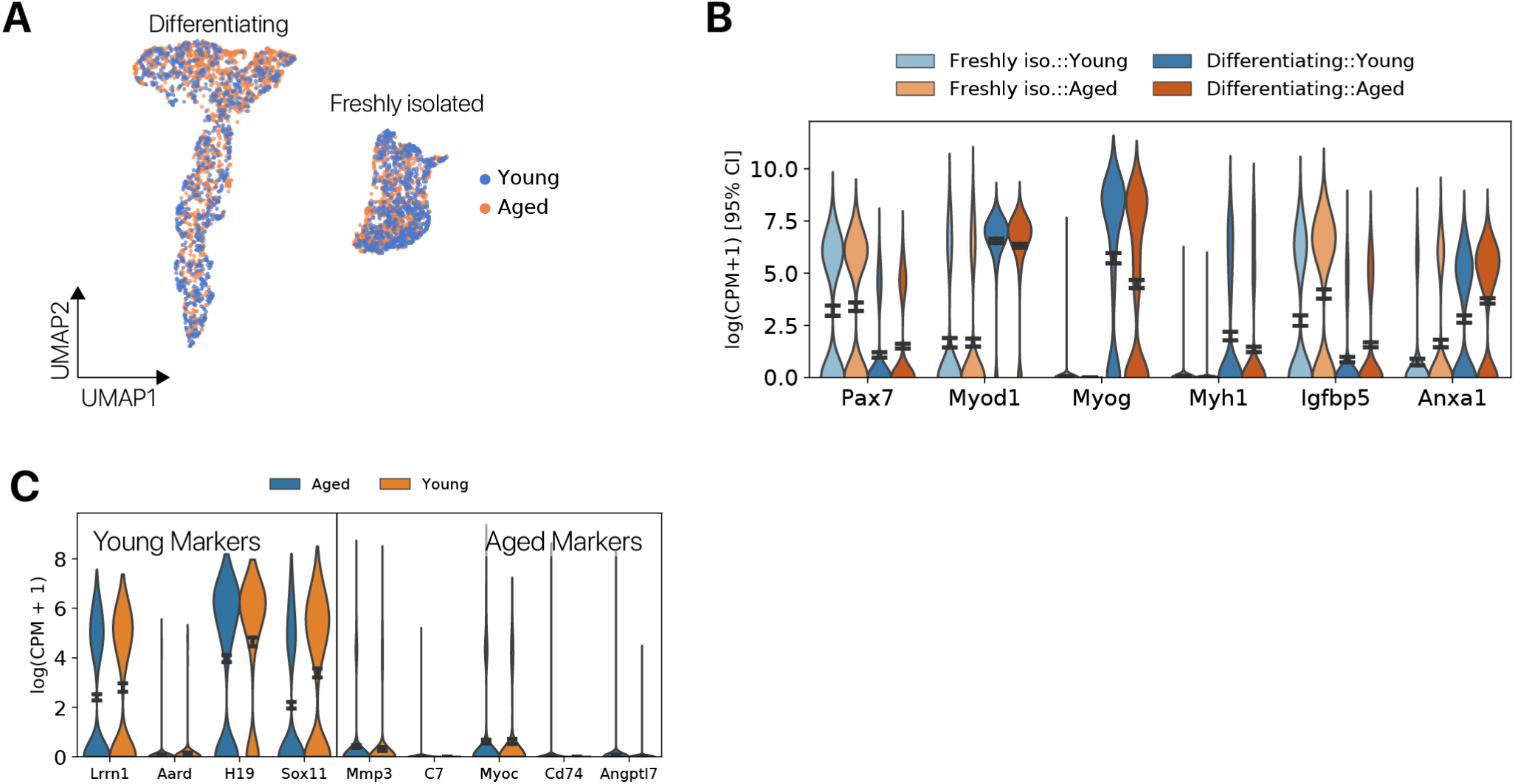
Differentially expressed genes with age in myogenic cells are consistent with previous reports. **(A)** Myogenic cells from the quiescent and differentiated time points presented in a UMAP projection. **(B)** Myogenic genes show expected expression changes with differentiation and aging markers show previously reported relationships with age. **(C)** Expression of the top 3 young cell markers and aged cell markers from a previous study of *in vitro* cultured MuSCs [51]. We find that some of the previously reported age markers repeat in our study, while others are undetected.

**Figure S6:**
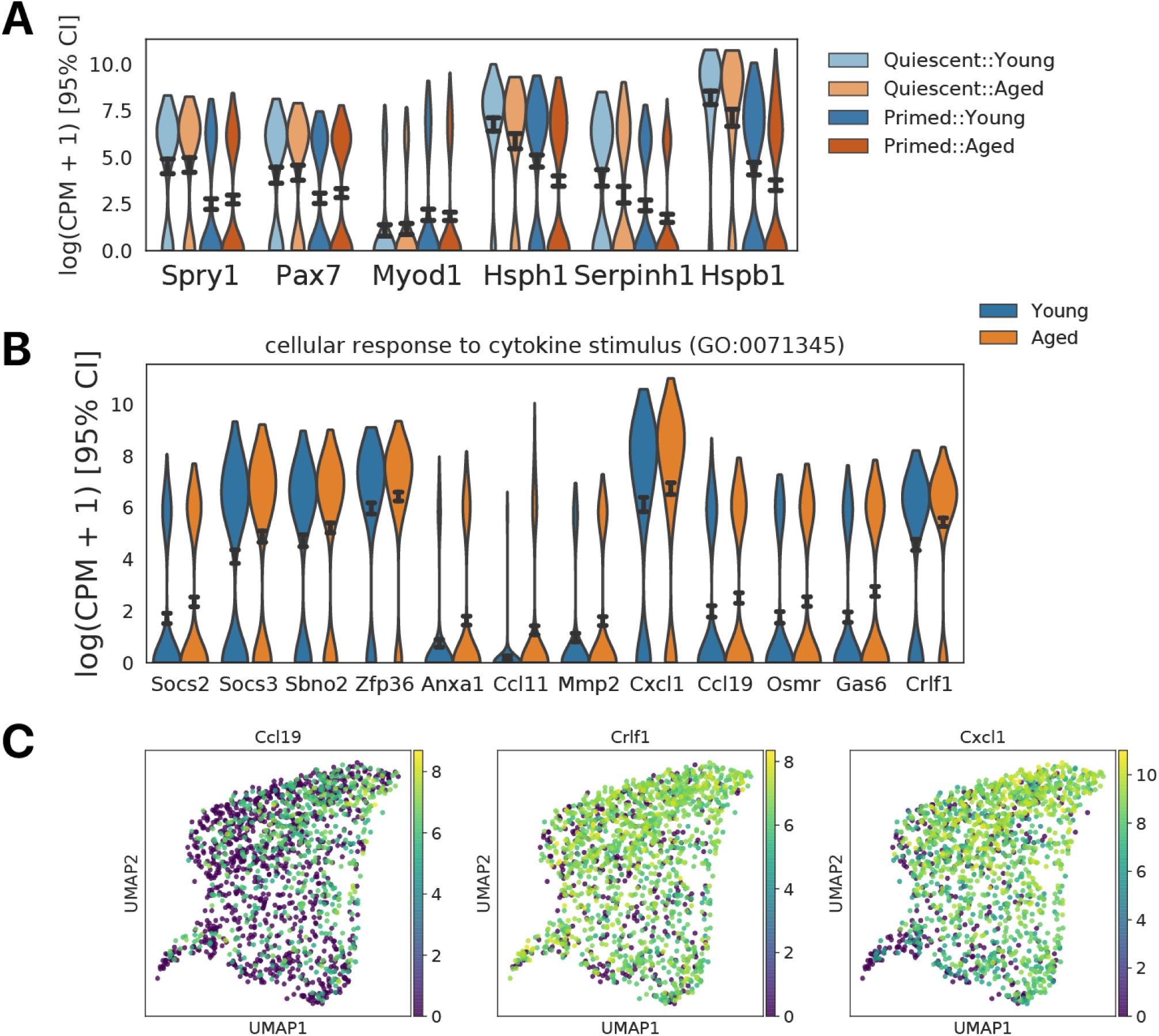
Aged MuSCs show increased expression of cytokine signaling pathways. **(A)** Violin plots of myogenic and UPR gene expression in quiescent and primed MuSC populations. The quiescent cluster is marked by *Spry1* and *Pax7*, and the primed cluster by *Myod1*. Expression of UPR genes is higher in the quiescent cluster. **(B)** Violin plots of cytokine stimulus genes in young and aged MuSCs. **(C)** Cytokine signaling associated genes displayed on a UMAP projection of MuSCs in transcriptional space. Cytokine signaling associated genes are expressed largely in the primed MuSC cluster.

**Figure S7:**
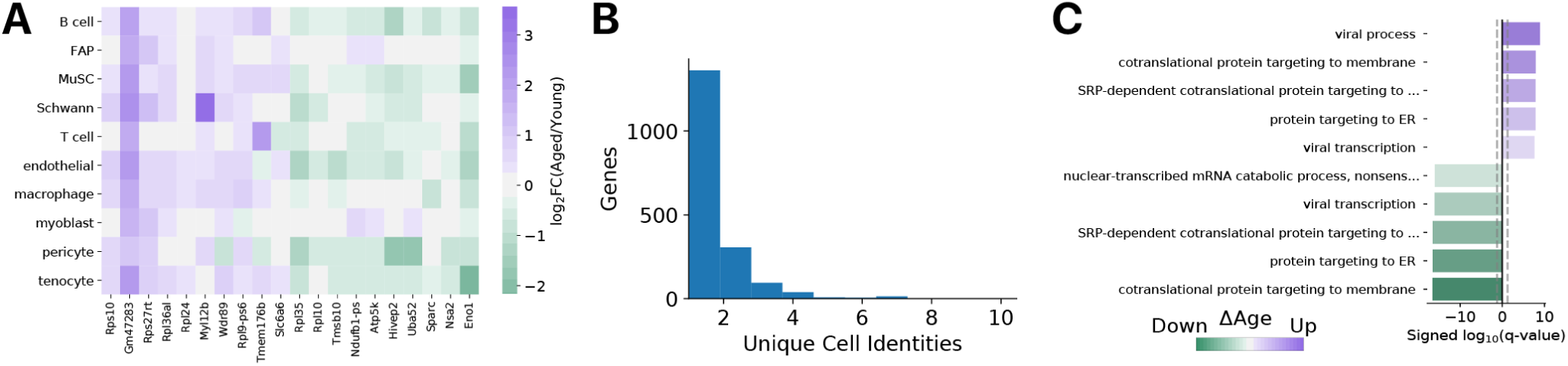
A subset of genes is commonly differentially expressed across cell types. **(A)** Heatmap of log_2_ fold-changes for common differentially expressed genes in each cell type. **(B)** Distribution of differentially expressed genes with age across the number of unique cell identities where those genes change in the same direction. Only a small subset of genes are changed in the same direction across many (*k* >= 3) cell identities. **(C)** Gene Ontology enrichment analysis of common differentially expressed genes reveals changes in SRP-dependent protein localization and co-translational protein targeting to the membrane with age.

**Figure S8:**
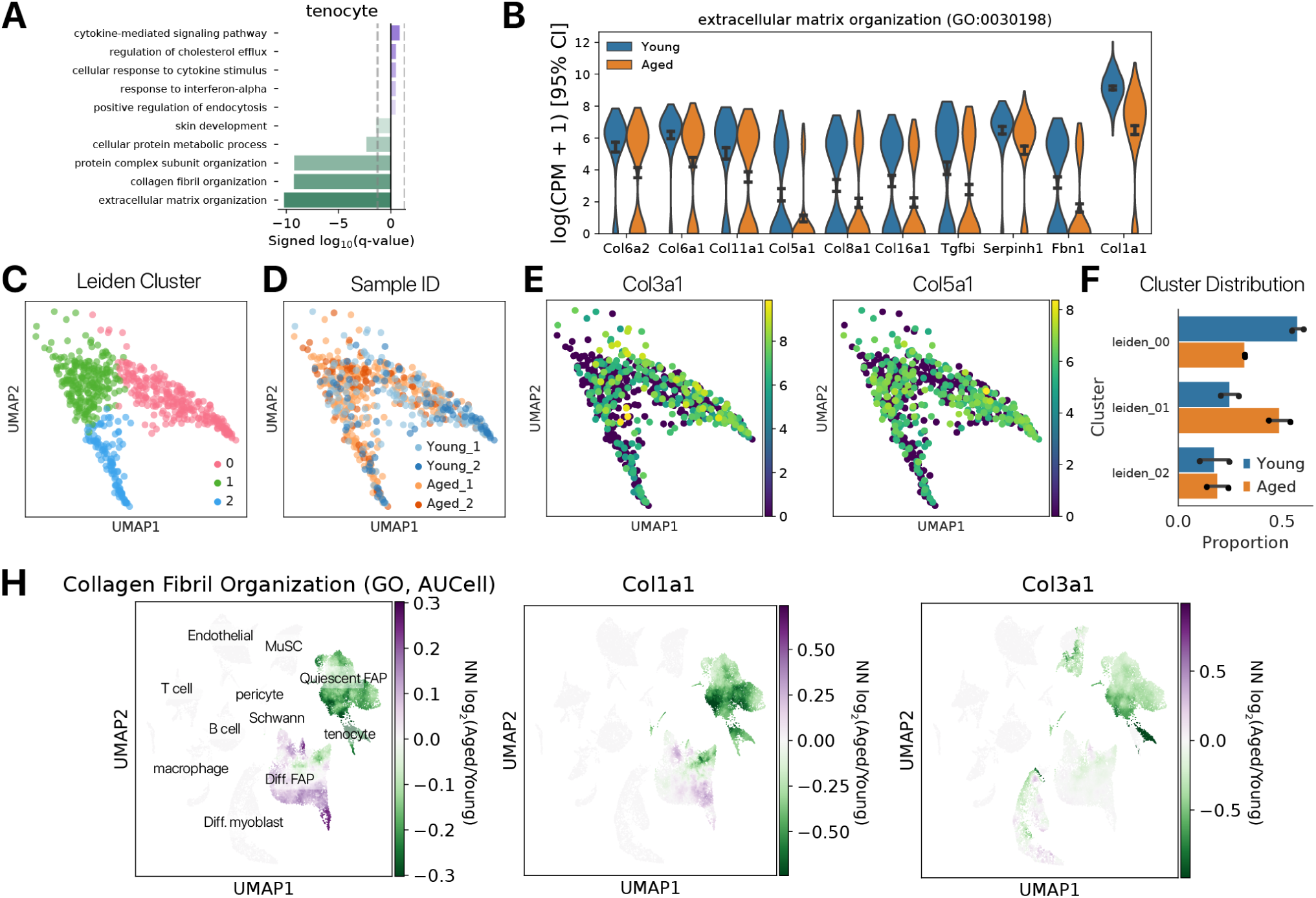
Aged tenocytes show reduced expression of collagen fibrils. **(A)** Gene ontology enrichment analysis for differentially expressed genes between aged and young quiescent tenocytes. **(B)** Violin plots showing the expression of a subset of genes driving enrichment of the “collagen fibril organization” GO term. **(C)** Leiden community detection identifies 3 subpopulations of quiescent tenocytes. Cluster labels are overlaid as colors on a UMAP projection of transcriptional space. **(D)** Sample IDs (age and animal replicate number) are overlaid as colors on a UMAP projection. We find that young tenocytes occupy a distinct region of transcriptional space (cluster 0) relative to aged tenocytes. **(E)** Expression of collagen genes is visualized on a UMAP projection, showing that cells in cluster 0 express higher levels of collagen genes than clusters 1 and 2. **(F)** Proportion of aged and young cells in each cluster. Cluster 0 is enriched for young cells, while cluster 1 is enriched for aged cells (*χ*^2^ test *p* < 0.01). Points represent the proportion of cells in a cluster for a single animal. **(H)** We computed Aged/Young fold-changes for the “collagen fibril organization” Gene Ontology gene set scored by AUCell in the local neighborhood of each cell (*k* = 100 nearest neighbors, Methods). We report a log_2_ fold-change of 0 for all local neighborhoods where few cells expressed the gene set. The collagen fibril organization gene set has decreased activity with age specifically in quiescent FAPs and tenocytes. Similar results are observed when scoring individual collagen genes (*Col1a1, Col3a1*), suggesting that FAPs and tenocytes are primarily responsible for changes in ECM gene expression with age.

**Figure S9:**
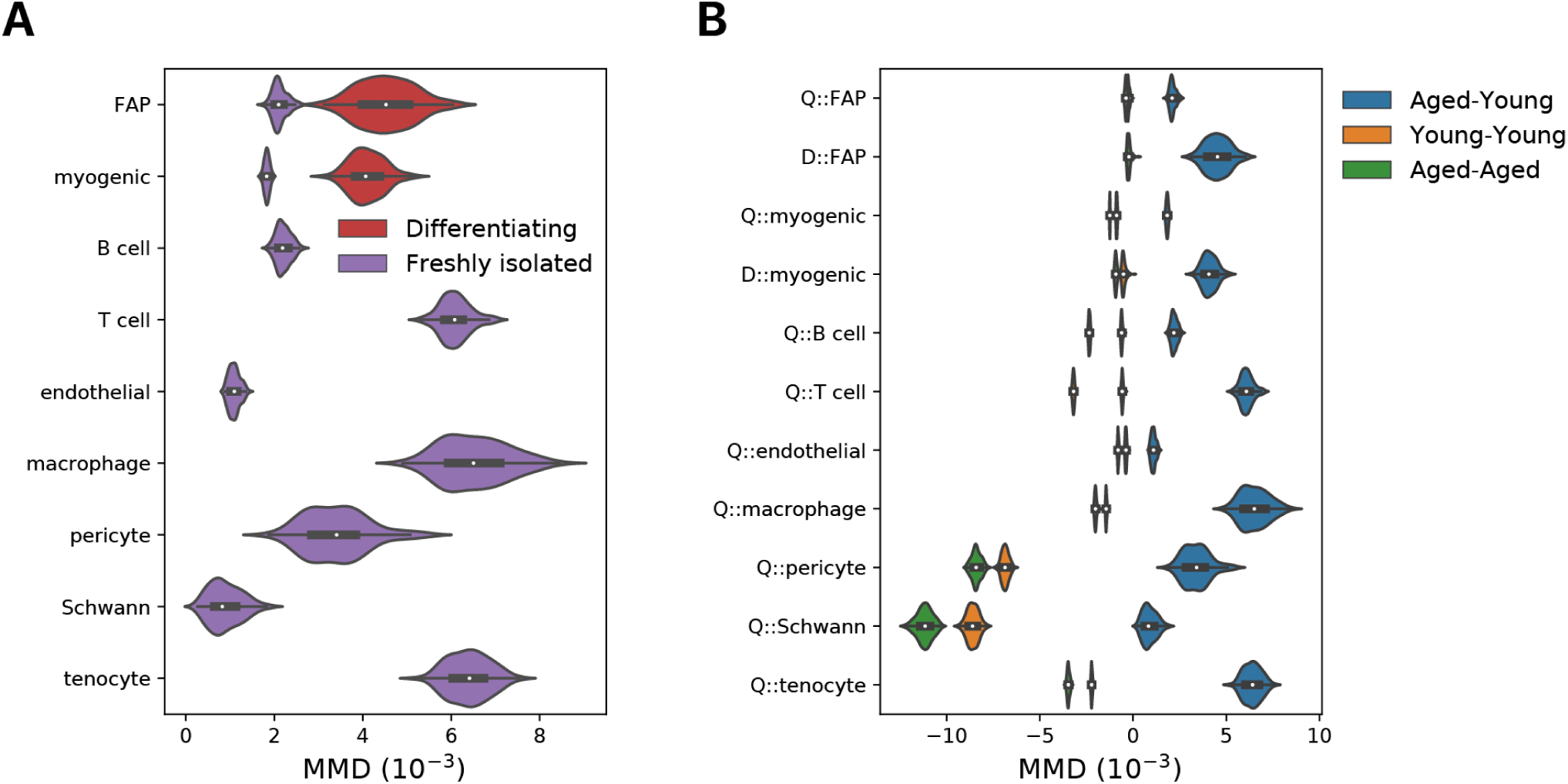
Aging magnitude estimated using maximum mean discrepancy distances in an NMF embedding. **(A)** Maximum mean discrepancy (MMD) distances computed between random samples of young and aged cells. The magnitude of aging increases with differentiation in both myogenic cells and FAPs (Wilcoxon rank sums, *p* < 0.001). **(B)** Heterochronic (Aged-Young) and isochronic (Young-Young, Aged-Aged) MMD distances computed using log-normalized counts. Isochronic MMD distances are all less than or near zero, as expected for distributions that are not significantly different (mean Kernel Two Sample Test *p* > 0.99).

**Figure S10:**
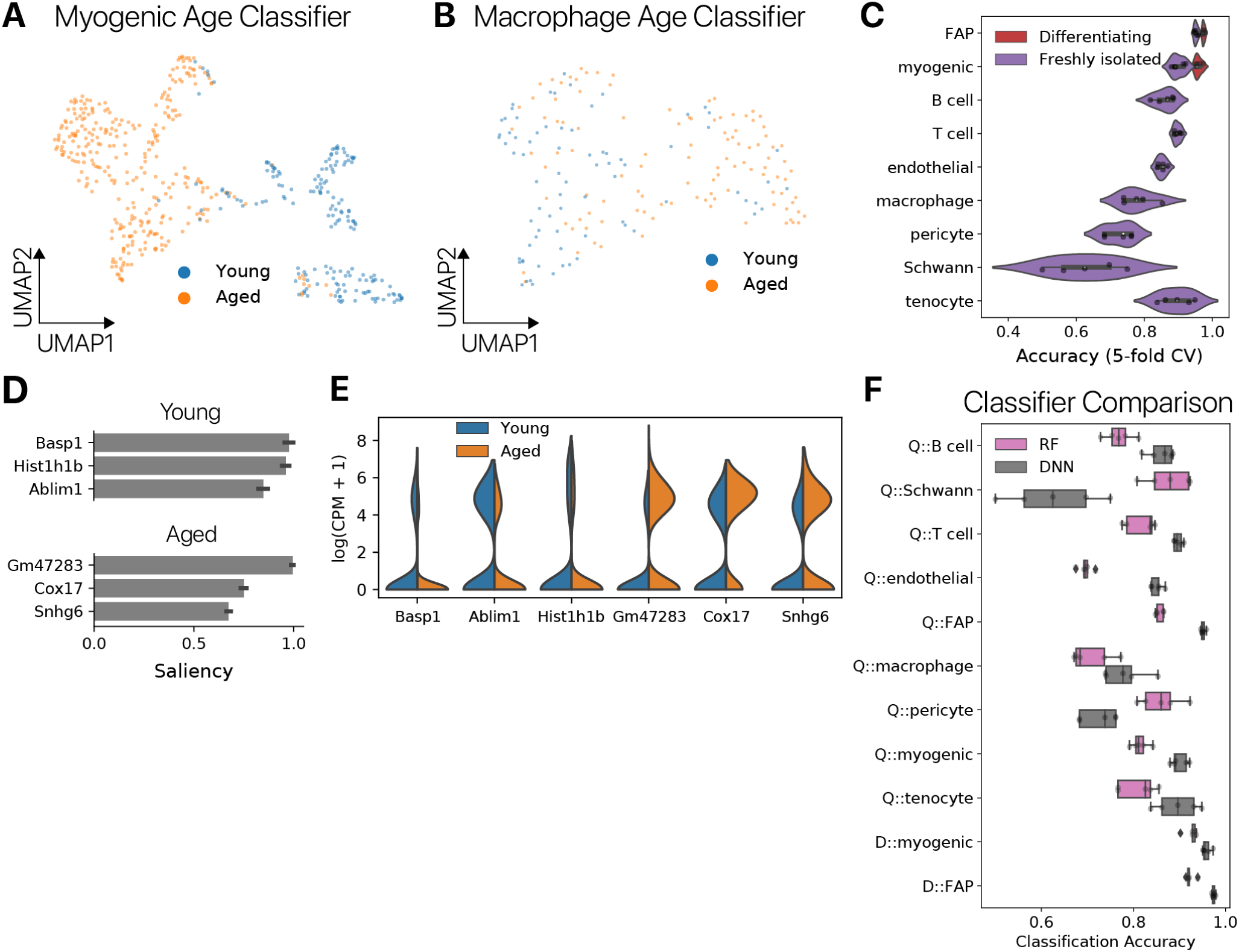
Young and aged cells exhibit distinct transcriptional states. **(A)** UMAP projections of embeddings from neural networks trained to classify young and aged myoblasts and **(B)** macrophages. Only cells from the held-out test set are shown. Macrophages show lower classification accuracy than myoblasts, and correspondingly show more overlap in the embedding. **(C)** Accuracy of deep neural network age classification models. Accuracy was assessed using 5-fold cross-validation, with separate data used for model selection and final evaluation. **(D)** Saliency analysis identifies important genes driving young and aged classification decisions in the myoblast age classifier. **(E)** Salient genes are differentially expressed across young and aged cells, consistent with intuitions. Not all salient genes show large fold-changes, indicating that dimensions which separate young and aged cell distributions do not always exhibit large mean shifts. **(F)** Comparison of age classification accuracies between deep neural network (DNN) models and random forest (RF) models. Neural network models show superior performance for most cell populations.

**Figure S11:**
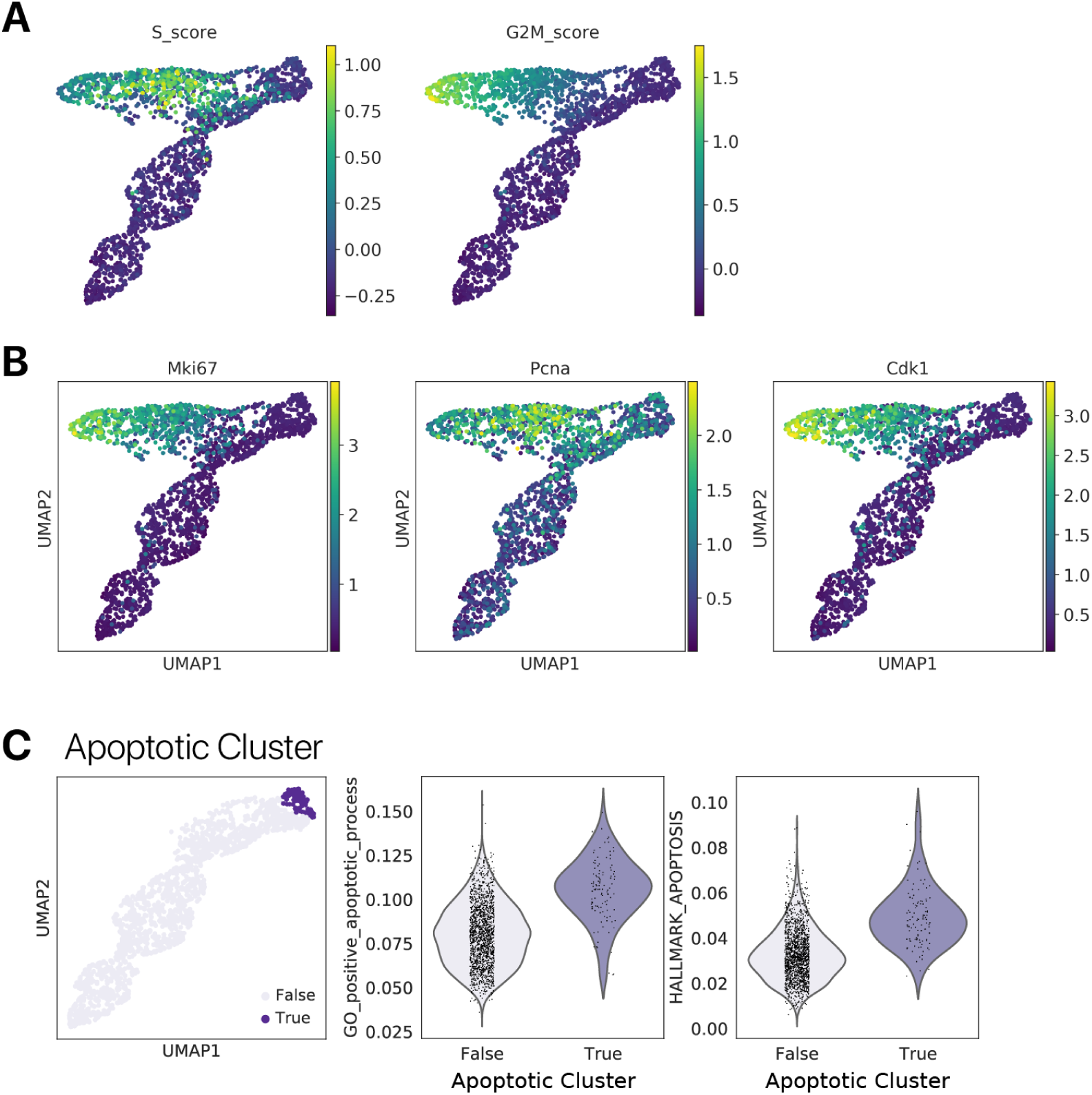
Identification of cycling and apoptotic differentiating myogenic cells. **(A)** Cell cycle gene set scoring identifies a cluster of cycling myogenic cells during differentiation. **(B)** Cell cycle marker genes likewise mark the same cluster of cycling myogenic cells. **(C)** A small cluster of cells at one end of the trajectory expresses significantly higher levels of apoptotic gene sets, as scored using AUCell. We removed these apoptotic cells from downstream analysis.

**Figure S12:**
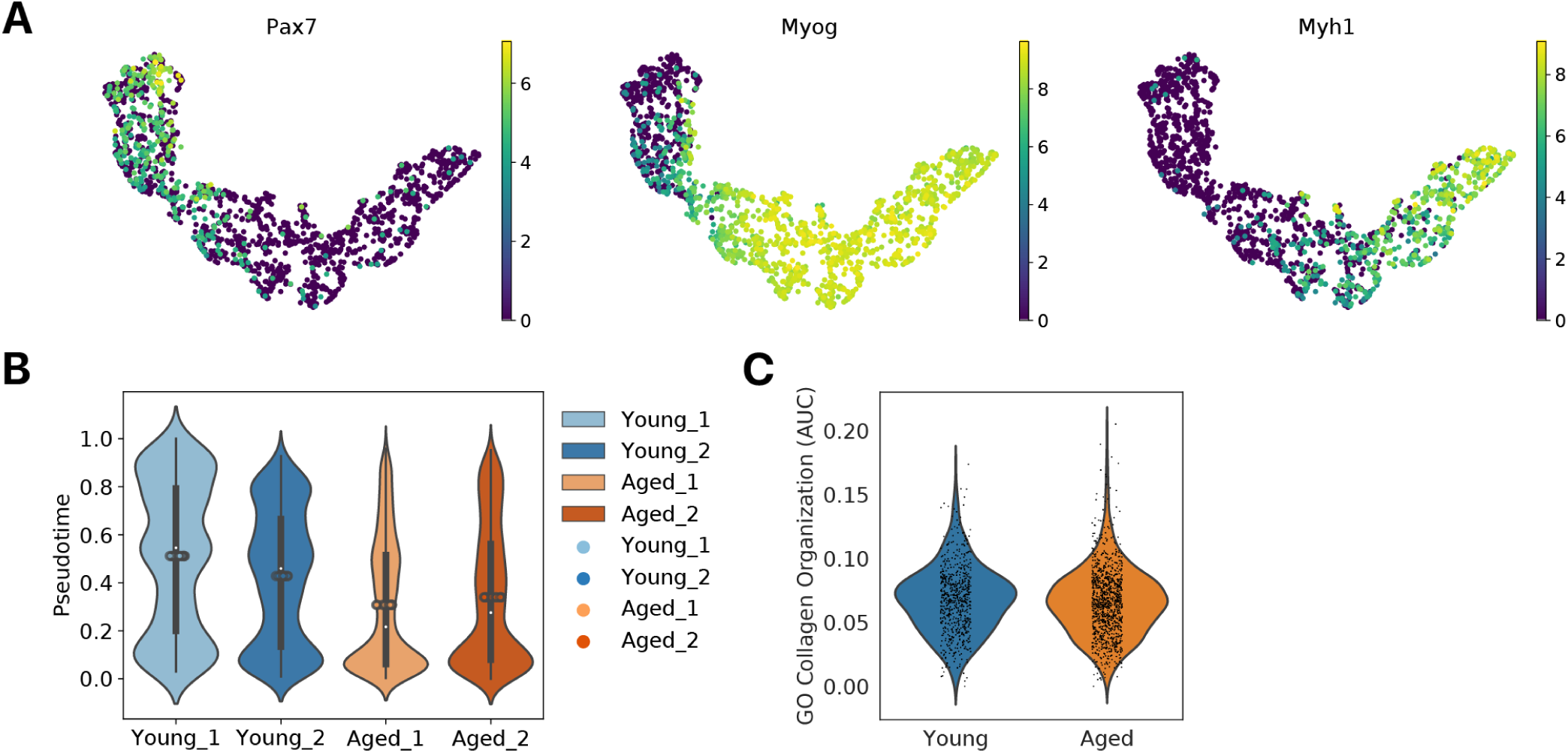
Young myogenic cells are enriched at later stages of the differentiation trajectory. **(A)** UMAP projection of differentiating myogenic cells displaying the expression of primitive cell marker *Pax7* and differentiated cell markers *Myog* and *Myh1*. **(B)** Violin plots of cell locations along a pseudotime trajectory for each animal of each age. Each violin contains points from 10 separate trajectories, each fit using a different root cell. Points plotted atop the violins are the mean pseudotime values for each animal for each of the 10 replicate trajectories. **(C)** Activity of the Gene Ontology (GO) Collagen Fibril Organization gene set scored using the AUCell approach. Aged myogenic cells do not show higher collagen gene set activity, suggesting that fibrogenic cell fates do not arise with age.

**Figure S13:**
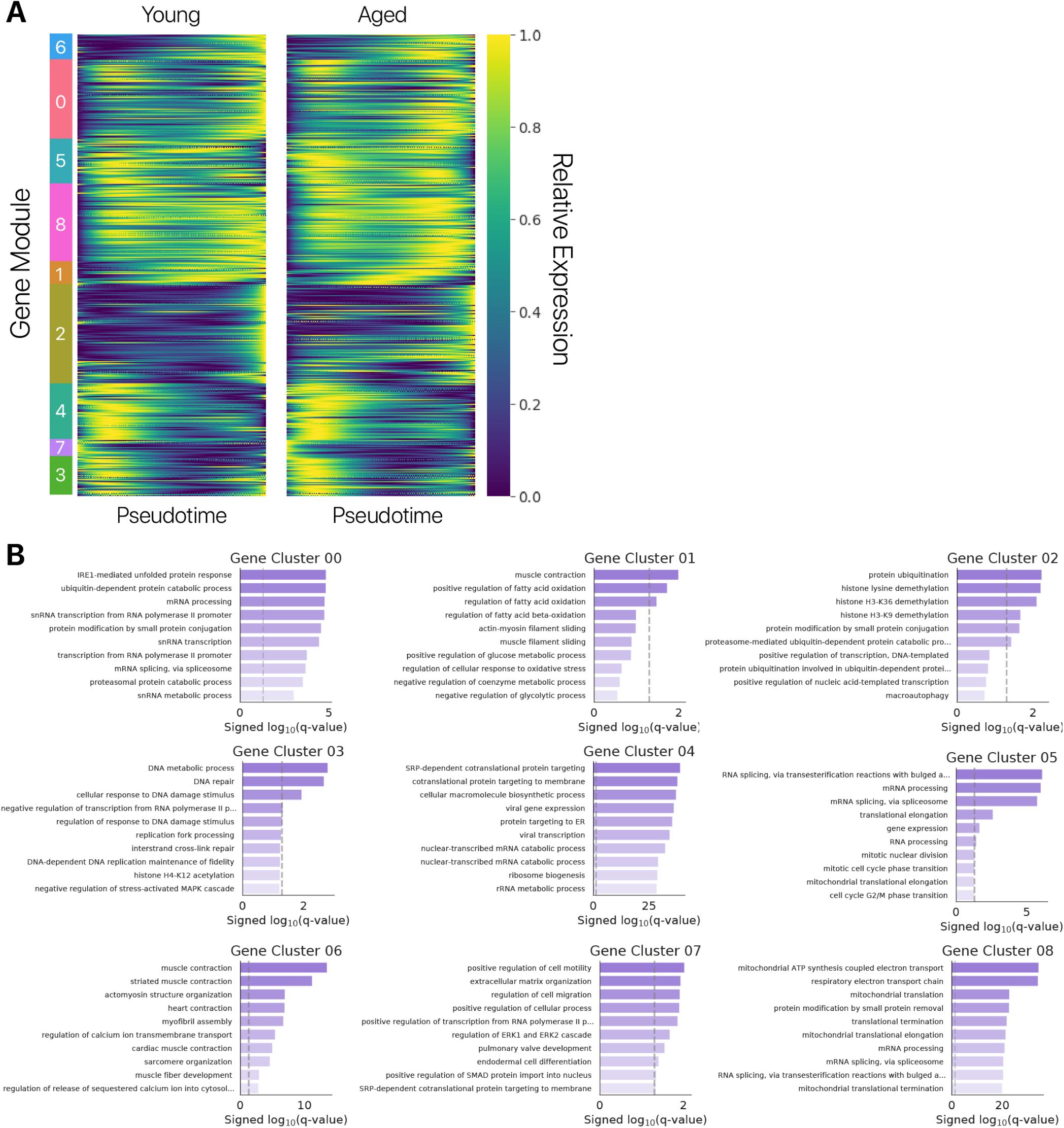
Pseudotemporal gene expression kinetics capture distinct modules of gene expression during myogenic differentiation. **(A)** Gene expression across pseudotime for young and aged cells, where each gene is a row and pseudotime increases across the x-axis. We identify 9 distinct gene modules that share similar kinetics during differentiation, including modules that decrease monotonically (3, 7), increase monotonically (2), or show non-monotonic behavior (5, 8). Young and aged cells display qualitatively similar gene expression kinetics across differentiation. **(B)** Gene Ontology enrichment results for each module in the myogenic differentiation program.

**Figure S14:**
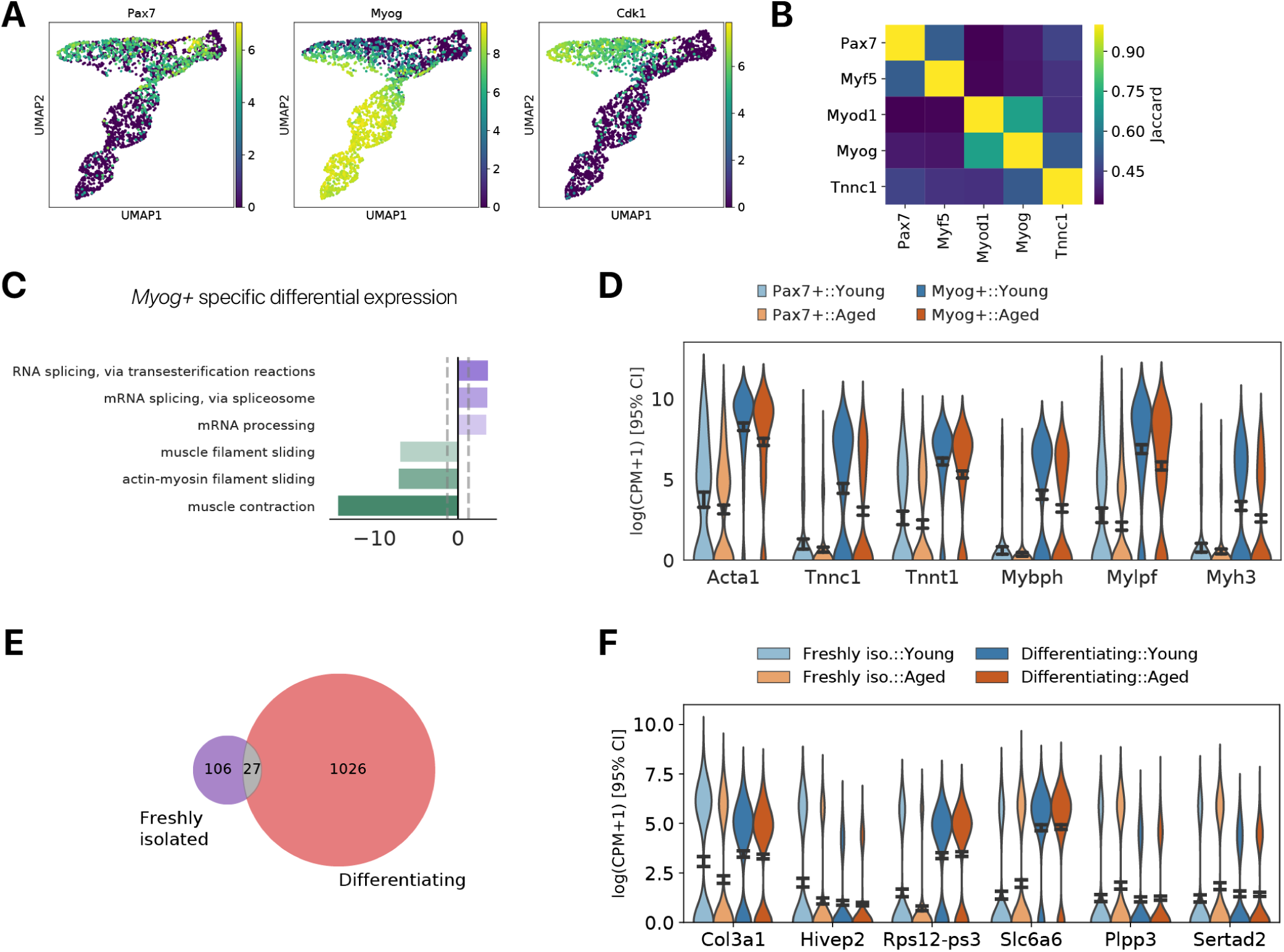
Myogenic differentiation masks and reveals age-related change. **(A)** Expression of *Pax7, Myog*, and *Cdk1* mark populations of myogenic progenitors, committed myoblasts, and cycling cells respectively. **(B)** Jaccard similarity metric (intersection over union) for differentially expressed genes detected in subpopulations of differentiating myogenic cells. We identified subsets based on the expression of known myogenic genes, indicated as labels on the matrix. **(C)** Gene Ontology enrichment analysis reveals that the “muscle contraction” term is more strongly downregulated in *Myog+* commited myoblasts than *Pax7+* progenitor cells. **(D)** *Myog+* cells show larger age-related changes in genes from the “muscle contraction” GO term than *Pax7+* cells. **(E)** Roughly half of all differentially expressed genes with age in quiescent MuSCs are no longer differentially expressed after differentiation, while differentiation manifests many new differentially expressed genes. **(F)** Expression of genes that are differentially expressed with age in quiescent MuSCs, but no longer differentially expressed after differentiation.

**Figure S15:**
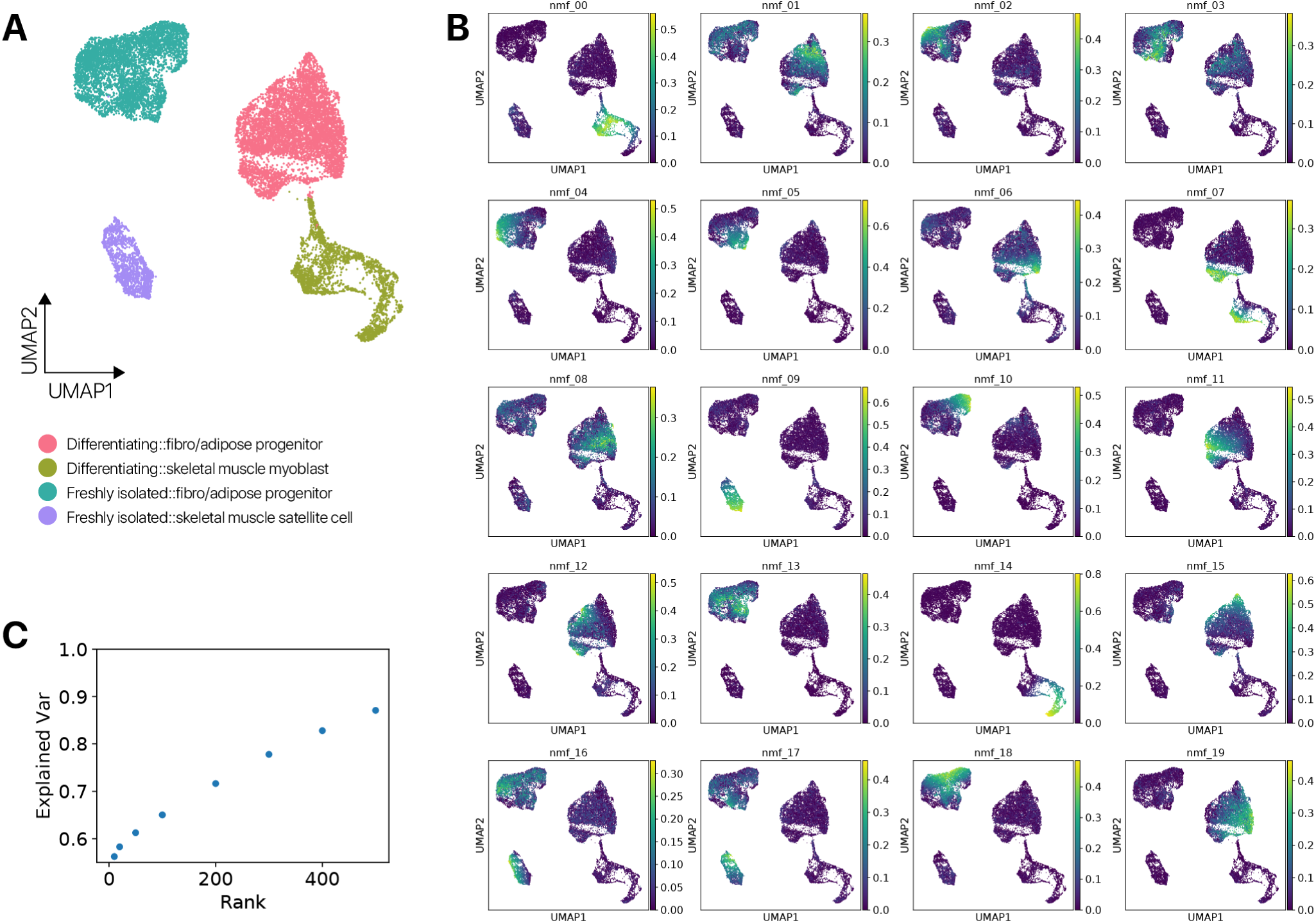
Non-negative matrix factorization identifies gene expression programs. We optimized a non-negative matrix factorization (NMF) embedding with 20 latent variables to capture gene expression programs in myogenic cells and FAPs. **(A)** UMAP projection of the NMF embedding. The NMF embedding preserves cell type and timepoint variation. **(B)** The activity of gene expression programs captured in the NMF latent variables is displayed in the UMAP projection. Some gene expression programs are primarily active in one cell type or timepoint, while others span cell types and timepoints. **(C)** Variance explained by NMF embeddings for a range of rank values *k*. The rank *k* = 20 embedding we select explains 58.3% of the variance in the data.

**Figure S16:**
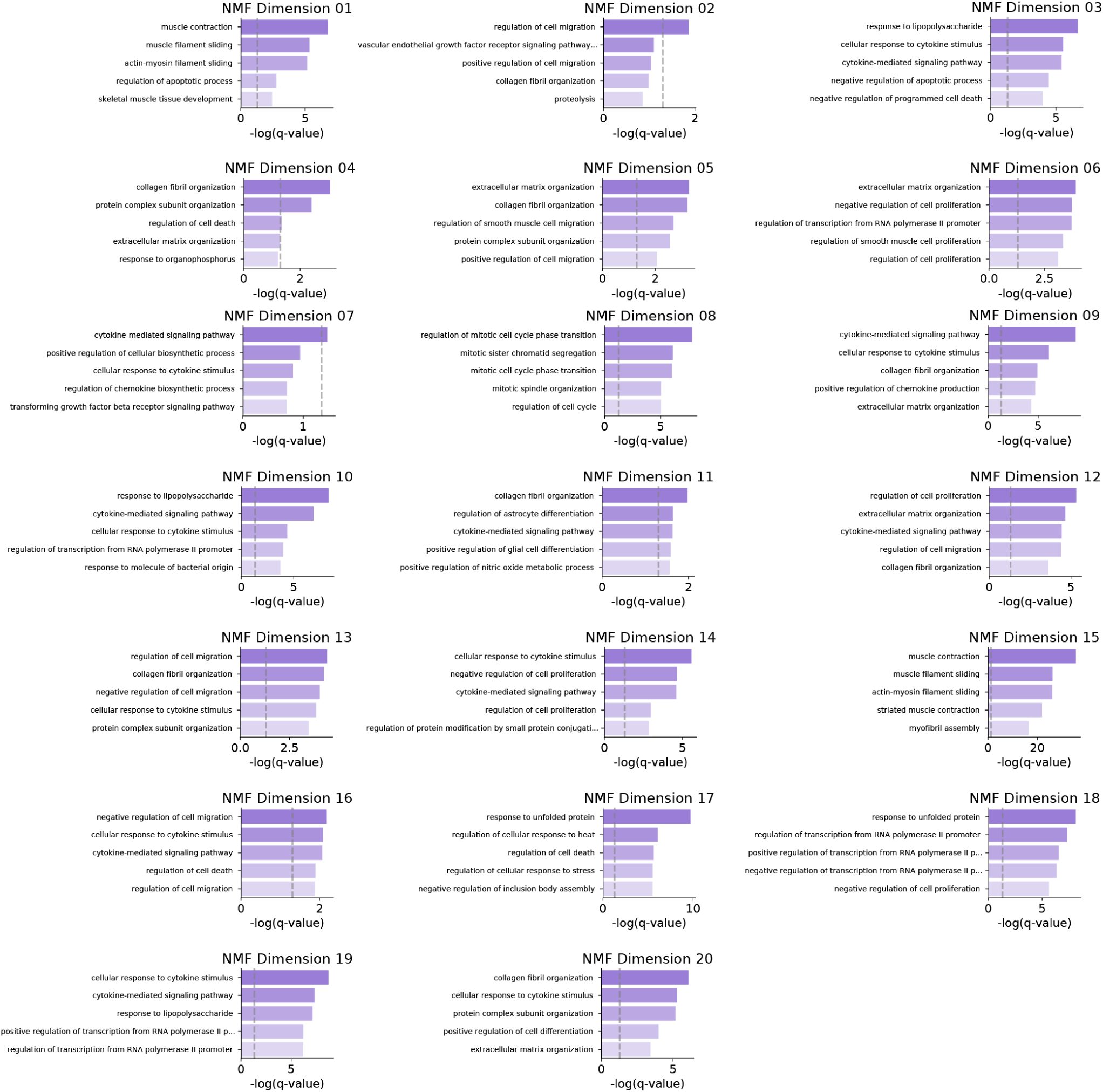
Gene Ontology analysis assigns semantic meaning to NMF latent variables. We identified genes associated with each NMF latent variable using Otsu’s threshold on the gene loadings matrix. We then performed Gene Ontology enrichment analysis to identify gene sets represented in each latent variable. From these enrichment results, we assigned a semantic name to each latent variable for downstream analysis and interpretation. The dashed line on the x-axis of Gene Ontology plots represents the *q* = 0.05 significance threshold.

**Figure S17:**
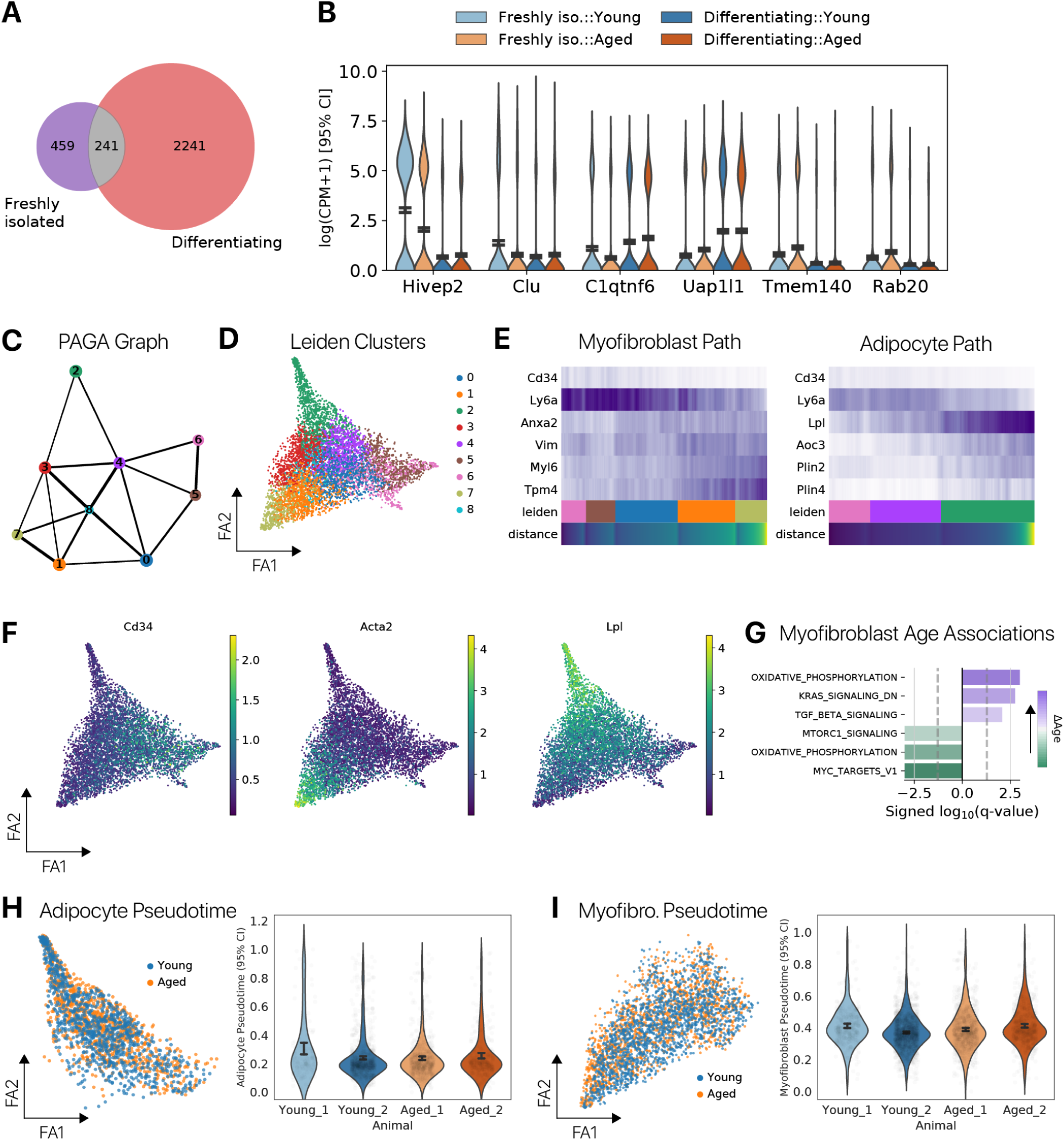
FAP differentiation masks and reveals age-related change. **(A)** Venn diagram of differentially expressed genes at the quiescent and differentiation timepoints. **(B)** Representative genes that are differentially expressed in one time point, and not in the other. **(C)** A coarse-grained topological graph showing relationships between clusters of differentiated FAPs. We constructed this coarse-grained graph with PAGA. **(D)** Leiden clusters of differentiated FAPs corresponding to nodes in the PAGA graph in **(C)**, projected in a force-directed graph layout embedding. **(E)** Expression of adipocyte or myofibroblast marker genes across the respective pseudotime trajectories. For each fate, we defined a path through the PAGA graph corresponding to the differentiation trajectory (Leiden row, colors correspond to PAGA nodes in **(C)**). Myofibroblast markers *Vim* and *Acta2* increase across myofibroblast pseudotime, while adipocyte markers *Lpl* and *Plin1/2* increase across adipocyte pseudotime. Progenitor markers *Ly6a* and *Cd34* decrease along both trajectories. **(F)** Marker gene expression for pre-adipocytes (*Cd34*), myofibroblasts (*Acta2*), and adipocytes (*Lpl*) presented in the graph layout embedding. **(G)** Genes upregulated with age in the myofibroblast trajectory are enriched for “TGF*β* signaling” genes (Hallmark, MSigDB, *q* < 0.05). **(H)** Distribution of young and aged cells in the adipocyte pseudotime trajectory and **(I)** myofibroblast pseudotime trajectory. There is no significant difference in the distribution of cells between ages (ANOVA *p* > 0.05) in either case.

**Figure S18:**
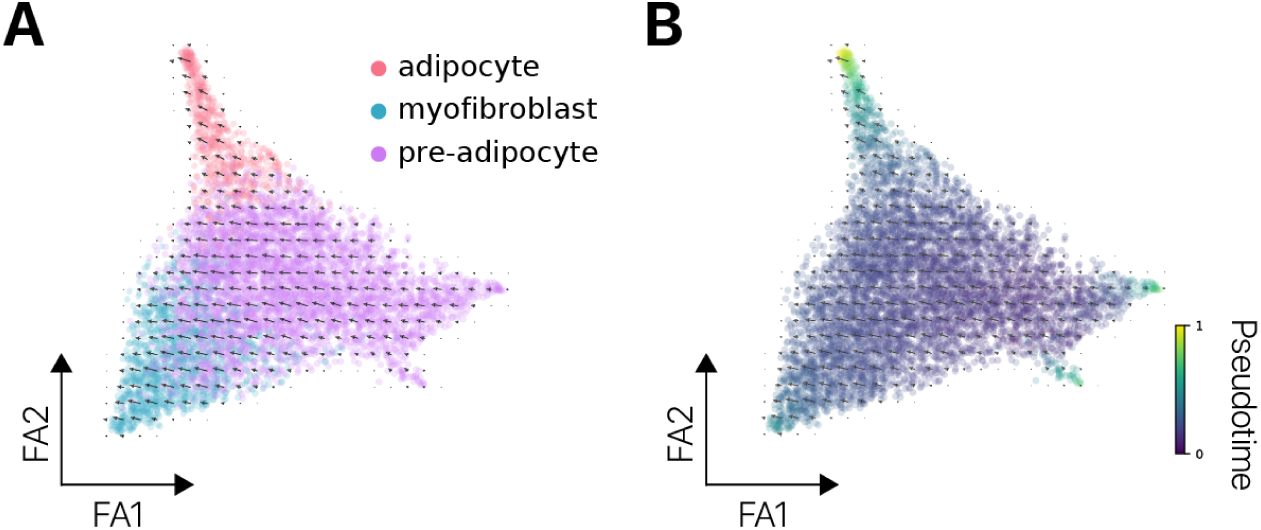
FAP pseudotime trajectories are consistent with RNA velocity estimates. **(A)** RNA velocity estimates are displayed as vectors atop a ForceAtlas2 graph layout embedding. We estimate velocity vectors in this embedding using a *k*-nearest neighbors regressor to project from PCA space to the ForceAtlas2 embedding coordinates (Methods). Cells are colored by fate assignment and **(B)** pseudotime coordinate.

**Figure S19:**
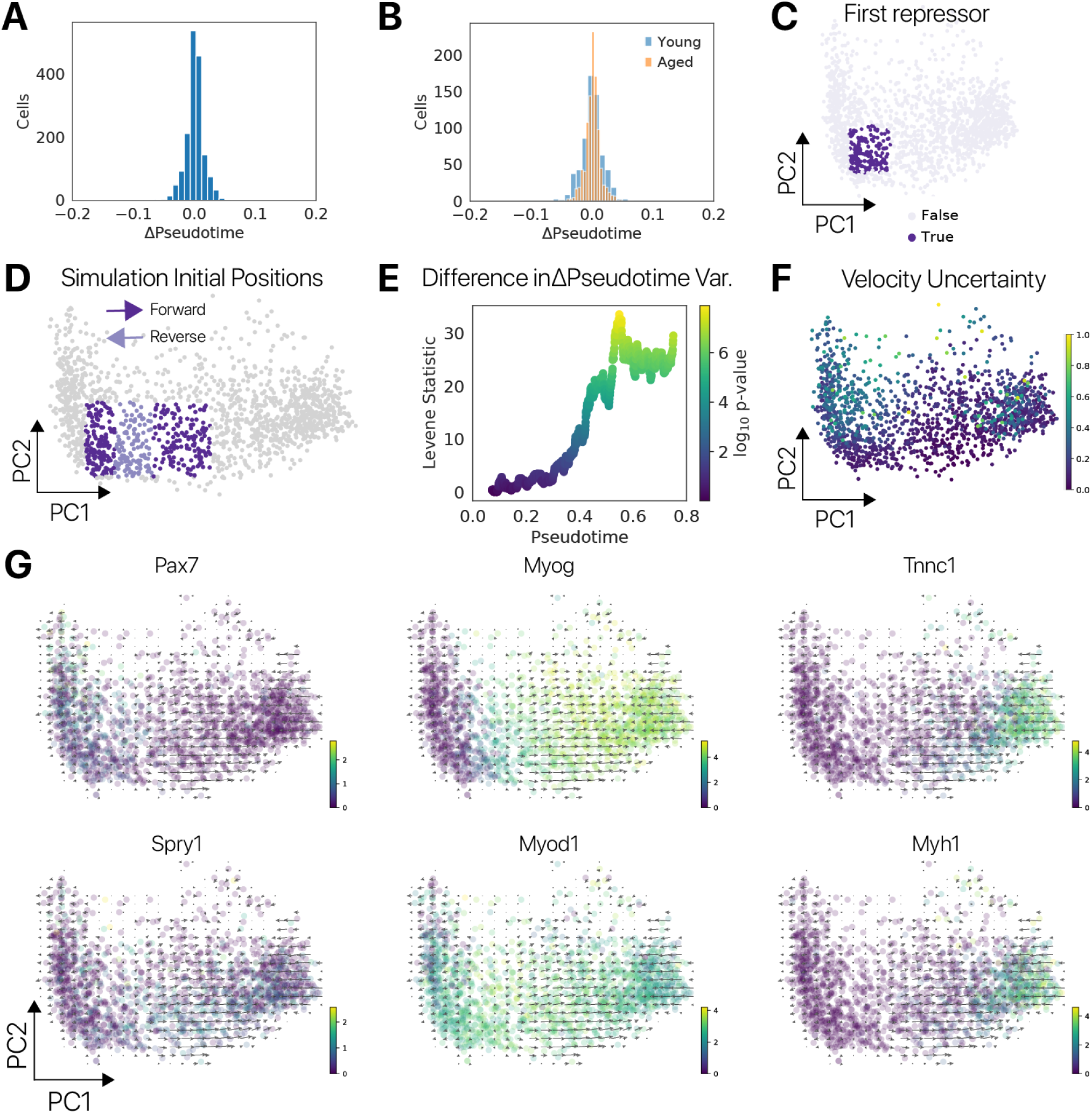
RNA velocity reveals a bifurcation and bidirectionality to myogenic differentiation. **(A)** Distribution of ΔPseudotime estimates for the differentiating myogenic cell population. ΔPseudotime is inferred by projecting each cell by its velocity estimate, inferring the pseudotime at the projected coordinate, and computing the difference from the observed pseudotime coordinate for that cell. **(B)** Distribution of ΔPseudotime values as in **(A)**, subset by age. **(C)** Location of the first repressor, highlighted in a PCA embedding of differentiating myogenic cells. **(D)** Initial positions for phase point simulations highlighted in a PCA embedding. **(E)** Levene’s test for difference in variance computed between young and aged ΔPseudotime values across the pseudotime trajectory. We found that the difference in variance in ΔPseudotime values between young and aged myogenic cells increases with differentiation. This suggests that differences in variance in the *Tnnc1+* attractor account for differences in ΔPseudotime variance observed in the whole population (as in **(B)**). **(F)** A metric of velocity uncertainty based on the correlation of velocity vectors in a local neighborhood is shown atop the PCA embedding (Methods). This metric identifies a region of low correlation (high uncertainty) in the *Tnnc1+* attractor, similar to our analysis of the variation in velocity vectors. **(G)** Expression of known myogenic regulators in the PCA embedding space with RNA velocity vectors visualized on top. We find that the bifurcation in velocity vectors localizes to the boundary of *Pax7* and *Myog* expression.

**Figure S20:**
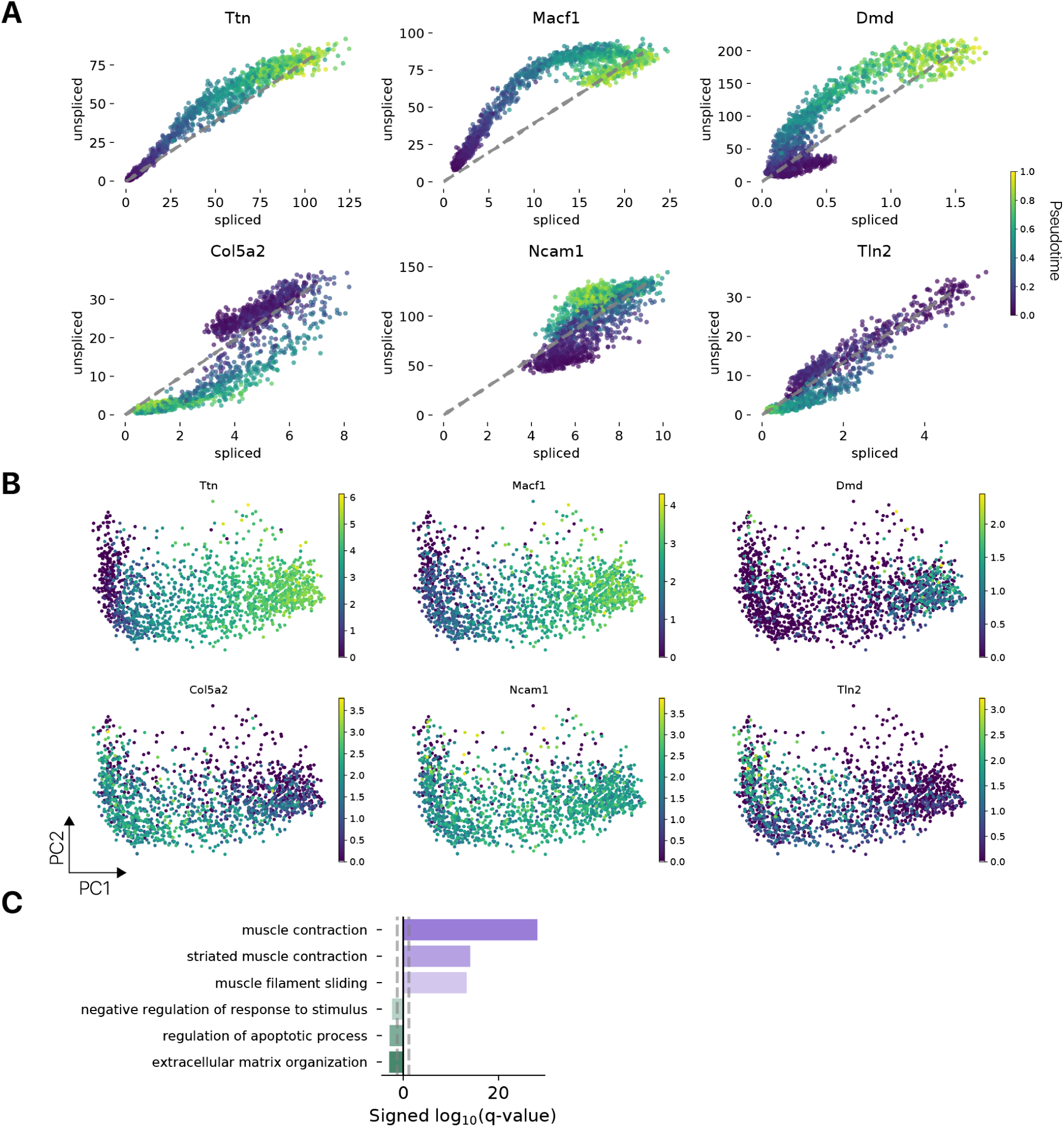
Myogenic genes, ECM, and stress response genes drive RNA velocity vectors. To identify genes that drive RNA velocity vectors in the PCA embedding, we computed the contribution of genes to the first principal component that is well aligned with pseudotime (Methods). **(A)** Spliced/unspliced phase plots for “forward” (top) and “reverse” (bottom) driver genes. Each point is a cell and the dotted lines represent the RNA velocity model fit. Myogenic genes *Ttn, Macf1*, and *Dmd* are induced in late pseudotime cells, driving a positive contribution in PC1 to the RNA velocity vectors. ECM gene *Col5a2* is induced in early pseudotime cells, driving a negative contribution to PC1 velocity vectors in these cells. In late pseudotime cells, repression of *Col5a2* (cells below model fit line) conversely drives a positive contribution to PC1 velocity vectors. *Tln2* displays similar dynamics, while *Ncam1* induction and repression have the inverse relationship with PC1 contributions. **(B)** Driver gene expression (log CPM + 1) visualized in the PCA embedding. Genes where induction provides a positive contribution to PC1 are enriched late in pseudotime, while genes where repression provides a positive contribution are enriched early in the pseudotime trajectory. **(C)** Gene Ontology enrichment analysis of the top *k* = 100 forward and reverse driver genes indicates that myogenic genes provide positive PC1 contributions while ECM, apoptotic, and stimulus response genes provide negative PC1 contributions to RNA velocity vectors.

**Figure S21:**
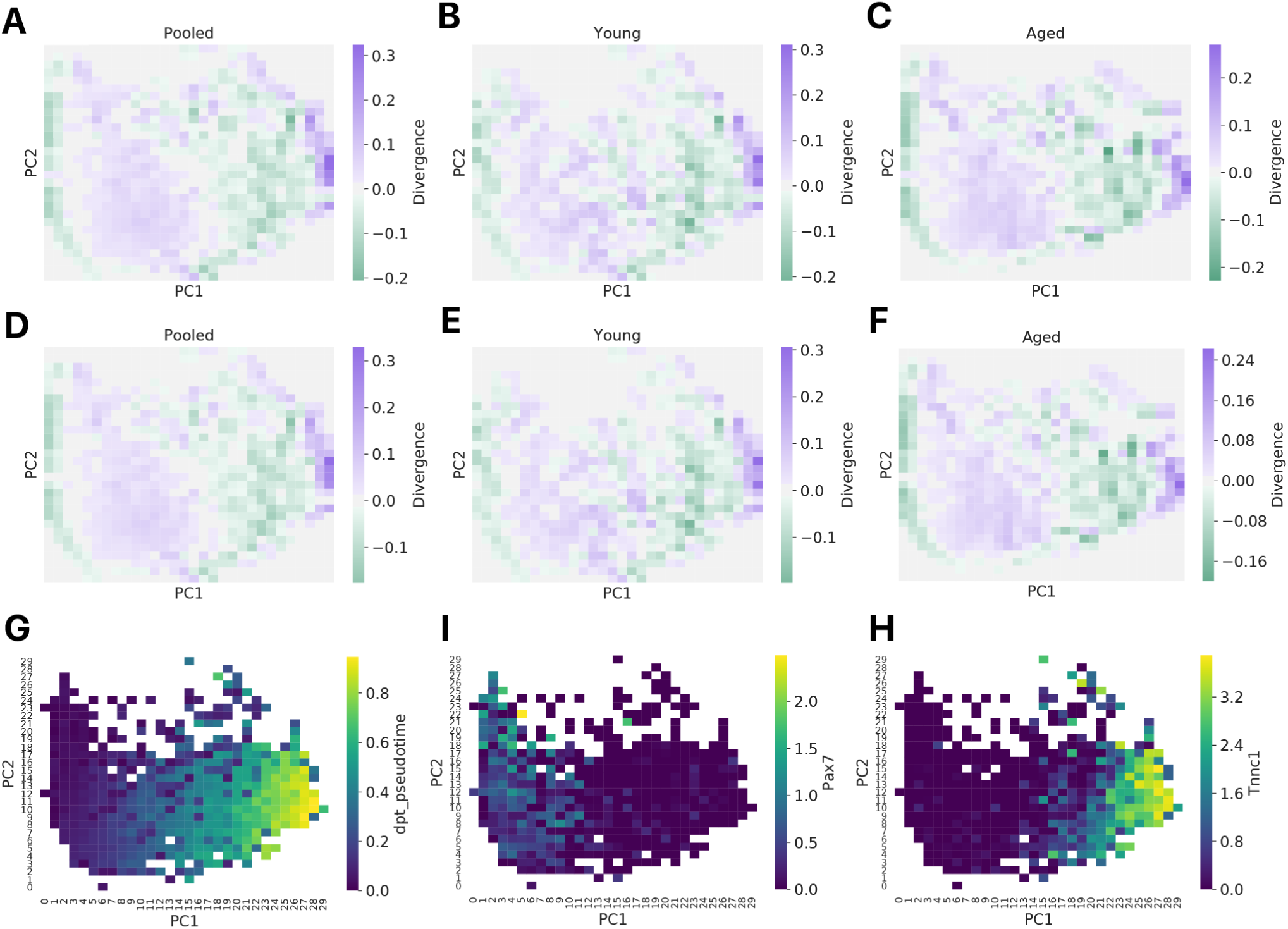
Divergence analysis reveals attractors and repulsors in myogenic differentiation. **(A)** Raw divergence maps computed from the coarse-grained velocity fields of all cells, **(B)** young cells only, or **(C)** aged cells only. We found few qualitative differences in the divergence computed for young and aged velocity fields. **(D)** Divergence maps computed as the mean of 100 bootstrapped samples from the distribution of all cells, **(E)** young cells only, or **(F)** aged cells only. For each bootstrap sample, we randomly selected 80% of the cells in the population. In these maps, we shrank the divergence value in a bin to 0 if the 95% confidence interval contains 0. These maps therefore show only divergence that is significantly different from 0. **(G)** Pseudotime overlaid in the coarse-grained PCA embedding. **(H)** Gene expression values (log CPM + 1) for *Pax7* and *Tnnc1* in the coarse-grained PCA embedding. The first attractor corresponds to *Pax7+* cells, while the second corresponds to *Tnnc1+* cells.

**Figure S22:**
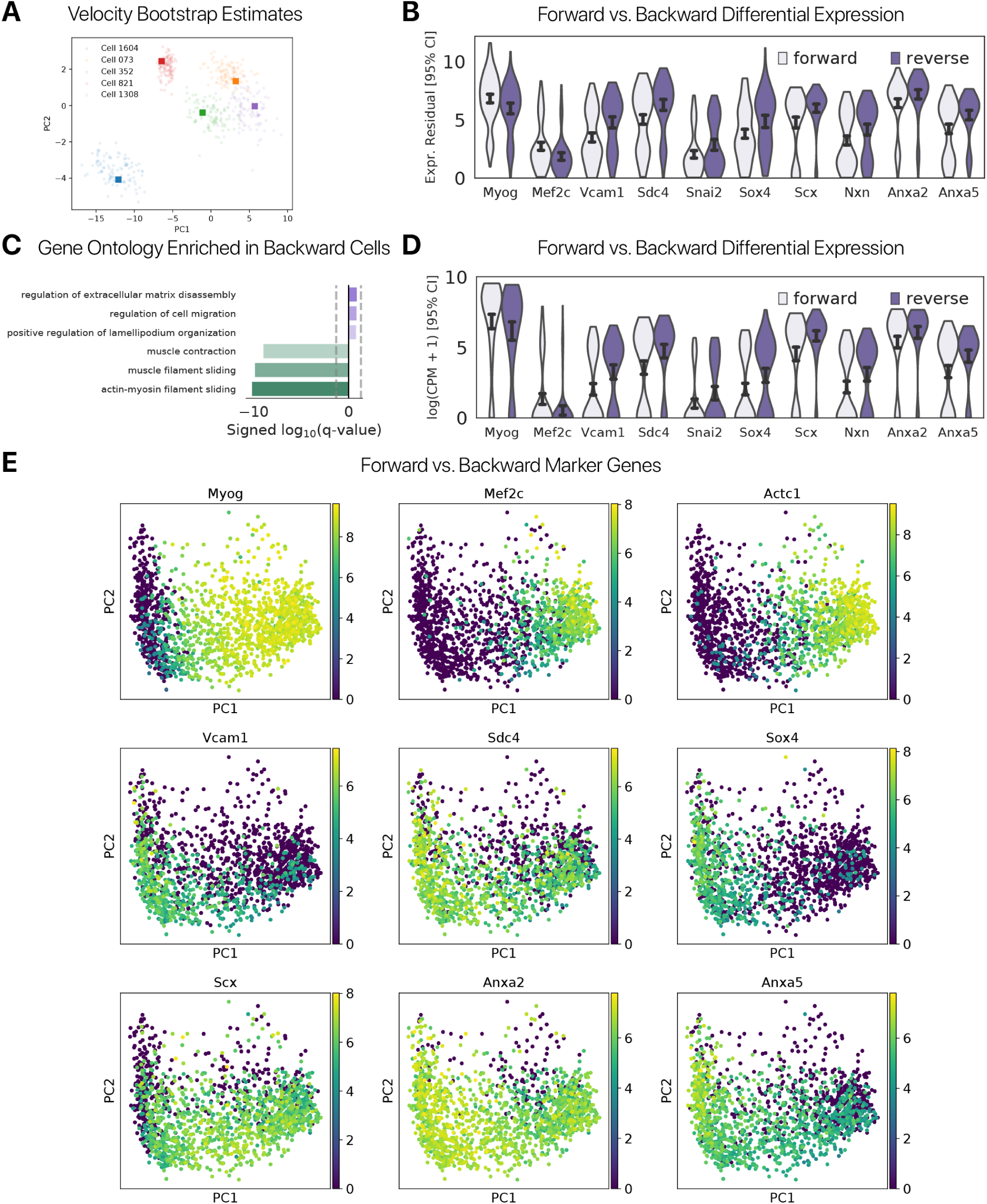
Differential expression reveals markers of myogenic lineage progression and reversal. **(A)** PCA projection of RNA velocity vectors derived using our molecular bootstrap sampling procedure. Colors indicate the cell of origin for each estimated vector. Circular points are estimates from a bootstrap sample, and square markers are velocity estimates computed with all observed reads. From the distribution of vectors, we derived RNA velocity confidence intervals. **(B)** Differentially expressed genes between cells moving forward (ΔPseudotime − 1.96*σ* > 0) and backward (ΔPseudotime + 1.96*σ* < 0) in the repulsor region. Values shown are residuals after regression of the pseudotime covariate from log(CPM + 1) counts and addition of minimum values to give a positive range. Myogenic transcription factors *Myog* and *Mef2c* mark cells progression forward, while primitive cell markers *Sdc4, Vcam1* and stress markers *Anxa2, Anxa5* mark cells moving backward. This suggests that relative levels of myogenic transcription factors, primitive cell markers, and stress signals may influence cellular decisions to move forward or backward through the differentiation program. **(C)** Gene Ontology enrichment analysis indicates that myogenic genes are downregulated in backward moving MuSCs relative to forward progressing MuSCs. **(D)** As in panel **(C)**, but with log(CPM + 1) values prior to regression of the pseudotime covariate. All markers maintain directionality in the absence of covariate regression.

**Figure S23:**
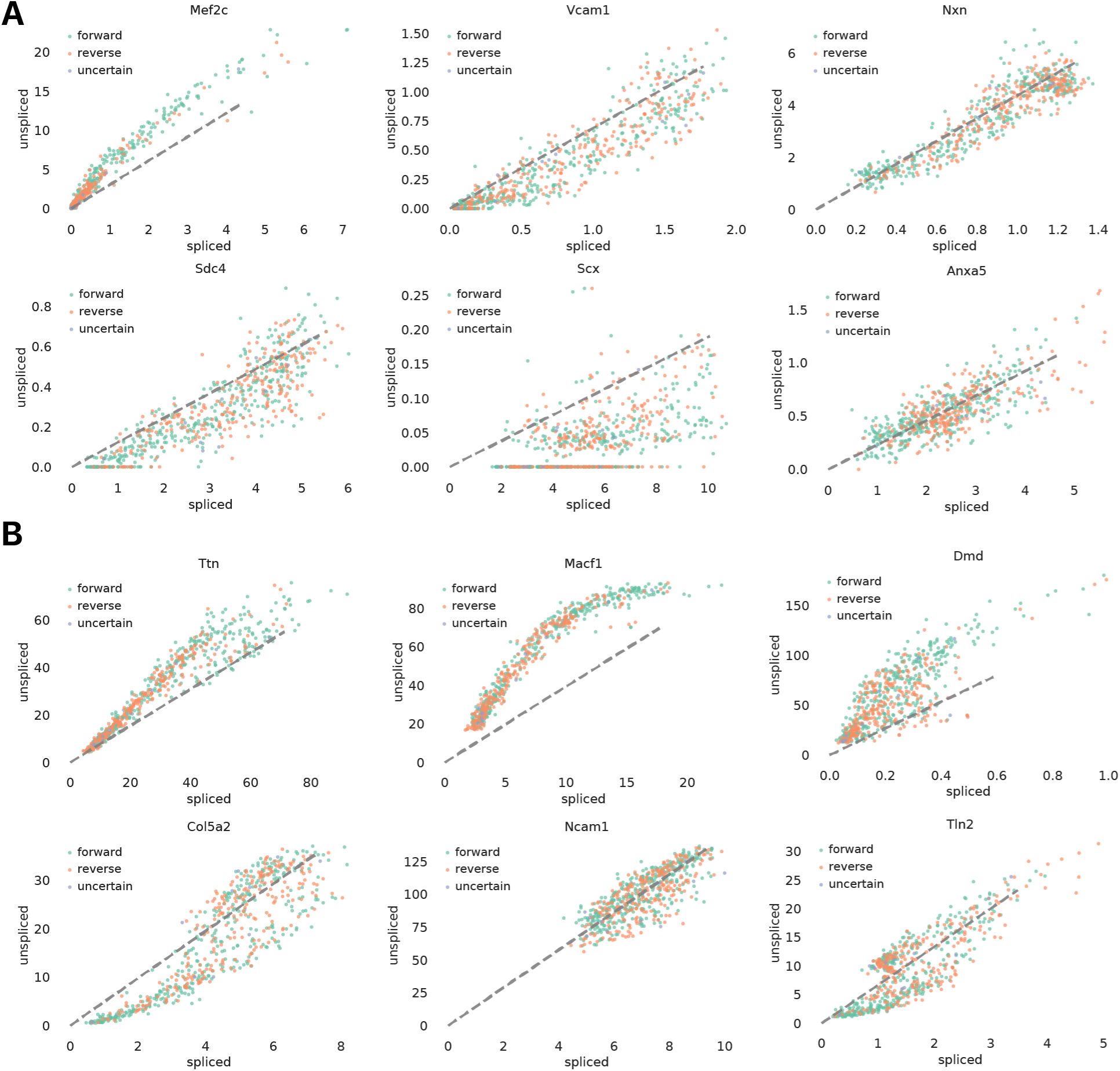
Differentially expressed genes between forward and backward moving cells indirectly influence velocity. **(A)** Spliced/unspliced phase plots differentially expressed genes between “forward” and “backward” moving cells. Each point is a cell and the dotted lines represent the RNA velocity model fit. Only cells in the repulsor region where we compute differential expression are shown. Points are colored by the direction of ΔPseudotime, where “uncertain” refers to cells where the 95% confidence interval contains 0. Forward and reverse moving cells do not show obvious differences in their distribution in phase space for these genes, with the exception of *Mef2c. Sdc4, Scx*, and the remaining differentially expressed genes are not used in computation of the velocity vectors. **(B)** Phase plots as in **(A)**, but using velocity genes we identified (Methods). Forward moving cells show large residuals for forward driver genes (*Ttn, Macf1, Dmd*), while backward moving cells show smaller residuals. Forward moving cells show negative residuals for backward driver genes (*Col5a2, Tln2*), whereas backward moving cells show positive residuals or smaller negative residuals.

**Figure S24:**
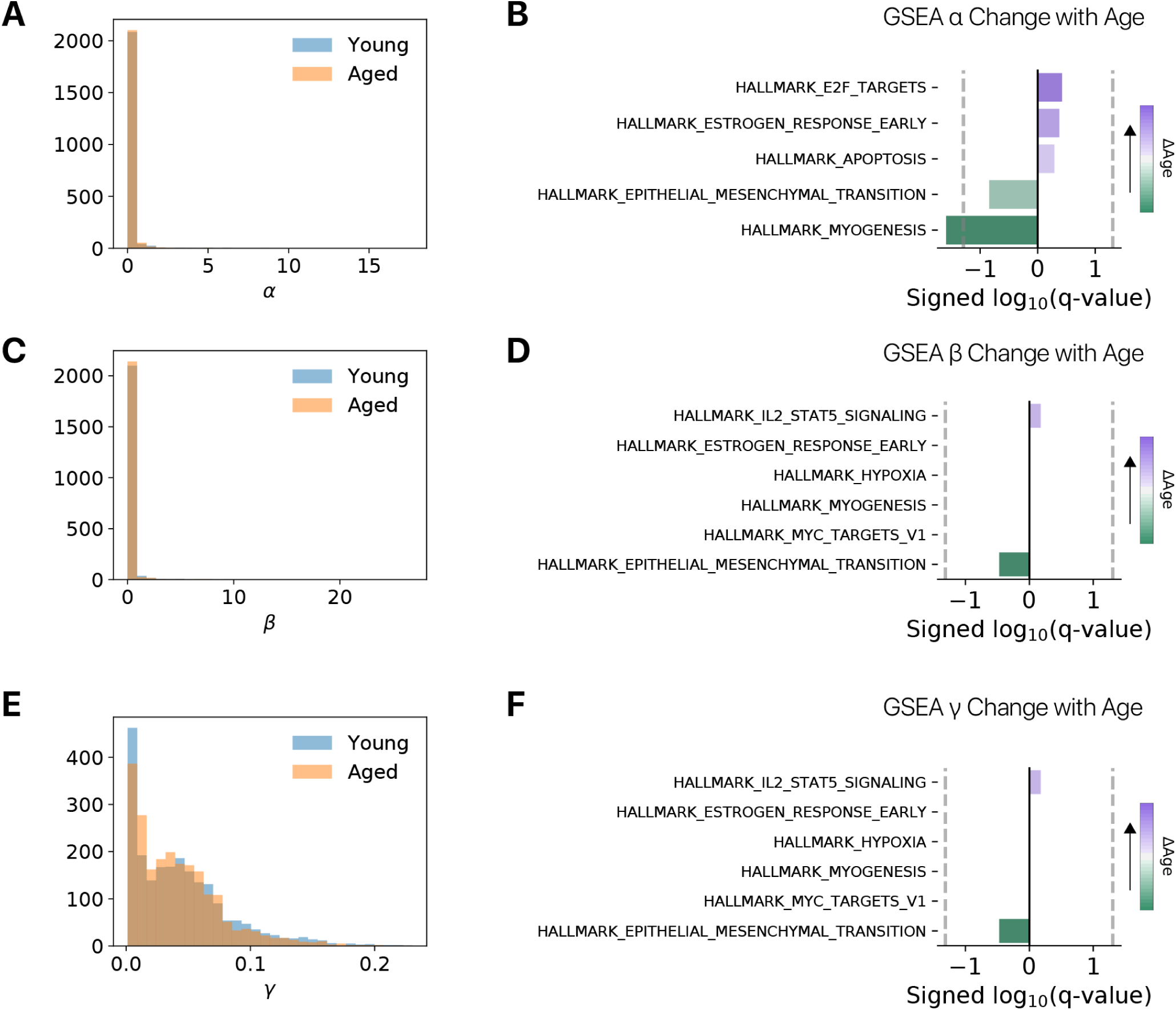
Inferred velocity parameters suggest aged myogenic cells transcribe myogenic genes at a lower rate. We fit separate `scvelo` dynamical models for young and aged myogenic cells to infer transcription (*α*), splicing (*β*), and degradation (*γ*) rates for each gene. We fit models to 100 bootstrap samples, each containing 80% of the cells from a given age. **(A)** Distribution of *α* transcription rate parameters across all bootstrap samples for young and aged myogenic cells. Distributions are similar across ages. **(B)** Gene set enrichment analysis (GSEA) reveals that aged myogenic cells have lower transcription rates *α* for genes in the “MYOGENESIS” Hallmark Gene Set. **(C)** Distribution of *β* splicing rate parameters across all bootstrap samples for young and aged myogenic cells. Distributions are similar across ages. **(D)** Changes in splicing rate parameters *β* across ages are not significantly enriched for any Hallmark Gene Sets by GSEA. **(E)** Distribution of *γ* degradation rate parameters across all bootstrap samples for young and aged myogenic cells. Distributions are similar across ages. **(F)** Changes in degradation rate parameters *γ* across ages are not significantly enriched for any Hallmark Gene Sets by GSEA.

## 7.1 Supplemental Tables

**Table S1:**
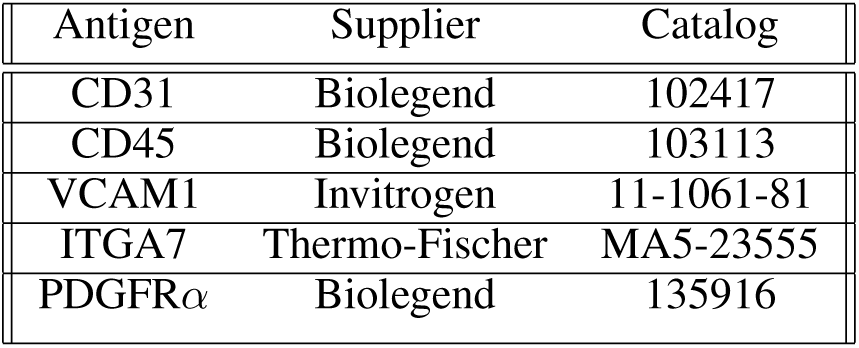
Antibody information.

| Antigen | Supplier | Catalog |
| --- | --- | --- |
| CD31 | Biolegend | 102417 |
| CD45 | Biolegend | 103113 |
| VCAM1 | Invitrogen | 11-1061-81 |
| ITGA7 | Thermo-Fischer | MA5-23555 |
| PDGFRα | Biolegend | 135916 |

